# Advancing Human iPSC-Derived Motor Neuron Models Using Glutamatergic Modulators for Synaptic Function Studies

**DOI:** 10.1101/2025.11.11.687934

**Authors:** Iman Rostami, Jana Leuenberger, Grischa Ott, Thomas Nevian, Benoit Zuber

## Abstract

Understanding human synapse structure and function at nanoscale resolution requires experimental systems that support both high-quality imaging and robust synaptogenesis. While brain tissue and primary neuronal cultures have been widely used, human induced pluripotent stem cells (iPSCs) offer key advantages: they enable patient-specific modeling and provide a renewable source of human neurons without relying on animal models. However, reproducible synapse formation in iPSC-derived neurons, particularly motor neurons (MNs), remains challenging due to variable differentiation efficiencies and low synaptic density. Here, we optimized MN differentiation to enhance synaptogenesis and enable high-resolution structural analysis using cryo-electron tomography (cryo-ET). Within 28 days, human iPSC-derived MNs developed into morphologically and functionally mature neuronal networks, characterized by phase-bright somata, elongated axons, and dense axodendritic branching. To further promote synaptic connectivity, we applied the glutamatergic modulators CX516 and CDPPB during differentiation. Their effects were validated by immunolabeling of synaptic markers, electrophysiology, calcium imaging, live-cell synaptic vesicle recycling assays, and ultrastructural imaging, including cryo-ET. This combined approach of optimized differentiation and targeted neuromodulation yielded reproducible MN networks with enhanced synaptic density and function, providing a robust in vitro platform for investigating human MN physiology, synaptic mechanisms, and disease-relevant synaptopathies.

## Introduction

In neuroscience, various immortalized cell lines have been employed as neuronal models, including human neuroblastoma (SH-SY5Y), mouse neuroblastoma (NB41A), catecholamine cells (PC12), embryonic neuronal precursors (LUHMES), and dopamine-containing hybrids (MN9D)^1–6^. Synaptogenesis, which results in interconnected neuronal networks, is a hallmark of neuronal development^7^. Although these models often display electrical excitability and generate action potentials, the formation of synapses with clearly defined ultrastructural features has not been conclusively demonstrated by electron microscopy (EM), and synaptic studies continue to rely predominantly on primary neuronal cultures^8^. Transmission electron microscopy, including cryo-electron tomography (cryo-ET), remains the gold standard for visualizing synaptic architecture at nanometer resolution. Unlike light microscopy techniques such as confocal imaging, EM resolves synaptic vesicles (SVs) and other ultrastructural features^9^. Primary neuronal cultures, often derived from rodent brain tissue, have therefore served as the standard in vitro model for studying synaptogenesis and network formation. These cultures reliably form synapses detectable by EM and have been instrumental in advancing our understanding of neuronal connectivity. However, they are animal-derived, which limits direct translation to human biology. Recently, human induced pluripotent stem cells (iPSCs) have emerged as a powerful alternative for generating neural lineages and glial cells^10,11^. The groundbreaking work of Takahashi and Yamanaka in 2006 demonstrated that introducing four transcription factors—Oct3/4, Sox2, Klf4, and c-Myc—into mouse embryonic or adult human fibroblasts could reprogram differentiated somatic cells into iPSCs^12^. This discovery revolutionized stem cell research, facilitating quick and reproducible generation of various cell types, including neurons^12,13^. Further studies have shown that patient-derived iPSCs represent a platform for developing and testing cell-based therapies^14–16^. In 2010, Hu et al. demonstrated that iPSCs could generate neuroepithelial cells and functional neurons similar to those produced by embryonic stem cells (ESCs) using the same transcriptional networks^17–19^. However, iPSCs exhibit lower efficiency and greater variability in differentiation, necessitating further improvements in differentiation protocols for potential use in cell transplantation^20,21^. Meanwhile, human ESCs face significant challenges, including ethical concerns related to the destruction of embryos and the risk of immune rejection^22^. These issues can be mitigated by using iPSCs, which provide a renewable, autologous source of cells in culture, making them an ideal candidate for regenerative medicine^23^. Today, dozens of clinical trials are underway using iPSCs, with neurological and cardiovascular diseases attracting attention^24^.

In this context, we present an optimized protocol for the maturation of iPSC-derived motor neurons (iMNs), which not only enables the formation of functional synapses but also facilitates their structural visualization using cryo-ET. iPSC-derived neurons generated using established differentiation protocols are capable of forming synapses; however, the efficiency and consistency of synaptogenesis remain inferior to primary neuronal cultures^25–28^. To promote synaptogenesis in vitro, we tested two neuromodulators with complementary glutamatergic mechanisms: CX516, an AMPAkine (a positive allosteric modulator of AMPA receptors), and CDPPB, a selective positive allosteric modulator of mGluR5^29–32^. AMPAkines prolong AMPA receptor channel opening and reduce desensitization, thereby enhancing excitatory transmission, which is typically associated with structural synapse strengthening. Notably, CX516 has been shown to restore altered single-channel kinetics of synaptic AMPA receptors in an Alzheimer’s disease animal model^33^. CDPPB facilitates mGluR5- and NMDA-dependent synaptic plasticity without inducing neurotoxicity or altering locomotor activity in rats^34^.Incorporating these modulators into our iMN differentiation protocol enhanced functional maturation and synaptic density, leading to measurable improvements in electrophysiological and ultrastructural indicators of synaptic connectivity.

## Results

### iPSCs Differentiate into Motor Neurons with Stepwise Maturation through Embryoid Bodies

Human iPSCs were successfully differentiated into MNs through an EB-based approach that recapitulated key stages of neural development^35^. The process involved three main steps: iPSC culture, EB formation, and MN differentiation (Figure 1). Initially, iPSCs were maintained for 17 days in six-well plates under standard conditions to preserve pluripotency, forming tightly packed colonies with clearly defined edges characteristic of undifferentiated cells. In the next phase, EBs were generated over 16 days by aggregating iPSCs into spherical clusters, which displayed smooth, well-defined borders by day 7. By day 16, neural progenitor cells formed and marked the onset of neural lineage commitment. Following their formation, EBs were dissociated into small clusters (1–5 cells) and seeded onto poly-L-ornithine and laminin-coated surfaces to initiate a 28-day MN differentiation protocol (Figure 1). By the first week after plating (DPD7), neural progenitors extended neurites, forming early networks. By the second week (DPD14), these networks became increasingly complex, with more branching and interconnectivity, reflecting advancing neuronal differentiation. By the third week (DPD21), cells exhibited distinct MN features, including large phase-bright soma and long axonal projections. By the final week (DPD28), cultures displayed a mature MN phenotype with dense, interconnected networks of axons and dendrites extending from well-defined cell bodies. The organized neuronal architecture was consistent with functionally mature neurons. These observations indicate stepwise differentiation of human iPSCs into iMNs, recapitulating key developmental stages, and yielding cells with morphological features of mature MNs.

**Figure 1.**
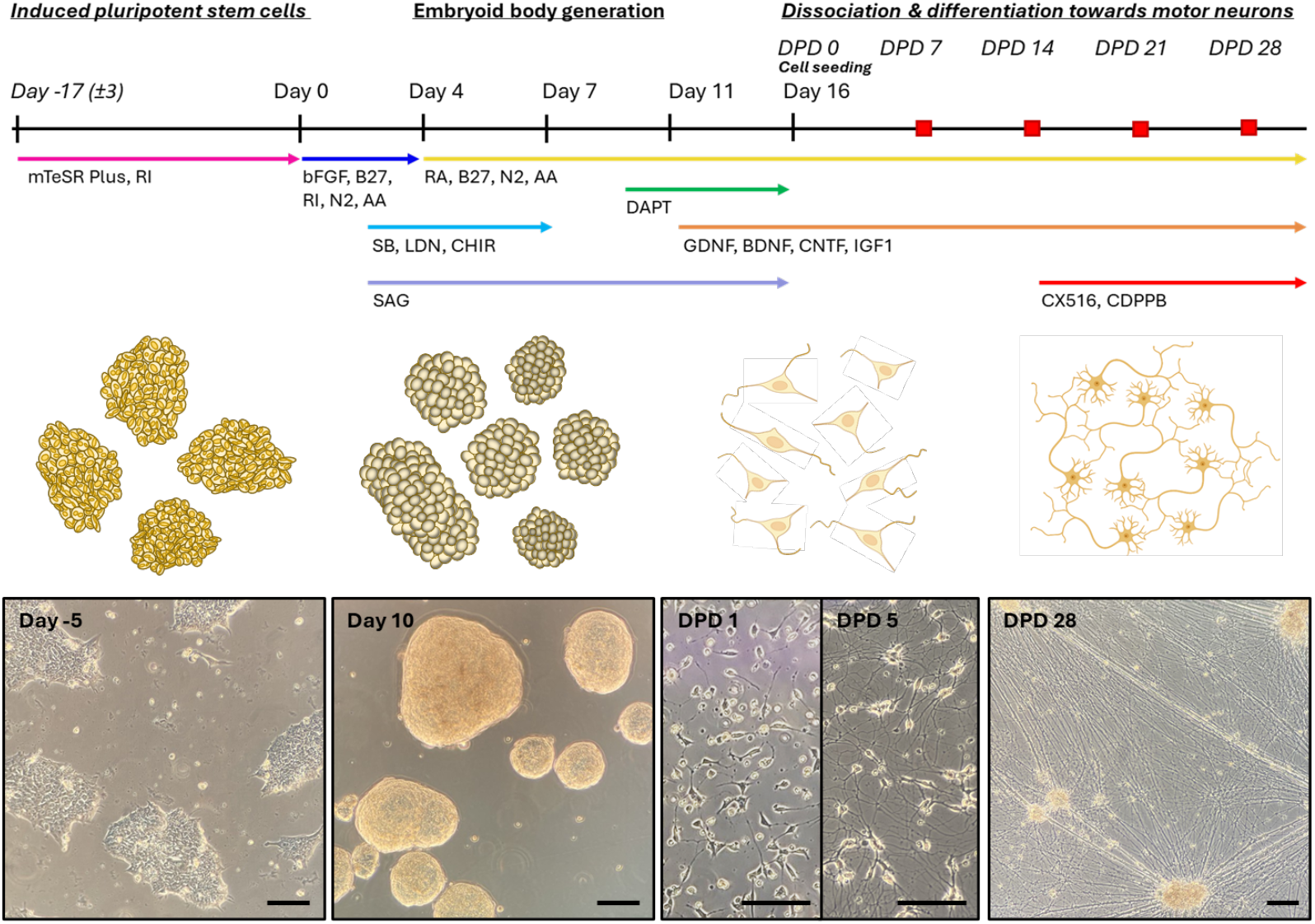
Differentiation timeline and morphological progression of iMNs. Schematic overview of the MN differentiation workffow from iPSCs, followed by representative phase-contrast images illustrating cellular morphology at key stages. The protocol consists of a multi-stage differentiation strategy. Neural induction is initiated at day 0 via EB formation, followed by patterning and MN specification using small molecules (e.g., RA, CHIR, and SAG) over the course of 16 days. Neuronal maturation continues post-single-cell plating (DPD: days post differentiation), with the addition of neurotrophic factors (e.g., BDNF, GDNF, CNTF, and IGF) supporting long-term survival and functional maturation. Neuromodulators are added at DPD14. Red squares indicate the time points at which analyses were conducted. Representative brightfield images at the bottom show the morphological progression of iMNs from Day −5 to DPD28. All images were acquired using a standard light microscope with a 10× objective. Scale bars: 100 μm.

The presence and localization of synaptic proteins were assessed by immunofluorescent staining (Figure 2, left panel). The entire neurite network was labeled with βIII-Tubulin, while dendrites were labeled by MAP2. Presynaptic puncta along neurites and cell body surface were labeled by synapsin-1 (SYN), synaptophysin (SYP), synaptic vesicle glycoprotein 2 (SV2) and Rab3A. Postsynaptic puncta along dendrites were labeled by PSD-95 (Figure S1). Close proximity or overlap of pre- and postsynaptic puncta (SYP/PSD-95) suggests that synapses were formed and can be used to quantify synaptic density and thus the extent of synaptic connectivity^36^. In confocal image stacks of iMNs at DPD28, we quantified an average of 83 ± 5.98 presynaptic puncta per 1000 µm^2^, of which 17.21 ± 3.52% overlapped with postsynaptic puncta, and 38 ± 10.59 postsynaptic puncta per 1000 µm^2^, of which 39.96 ± 11.33% overlapped with presynaptic puncta. These data, obtained from n = 5 fields, indicate an excess of presynaptic over postsynaptic structures in these cultures (Figure 3, box plot).

**Figure 2.**
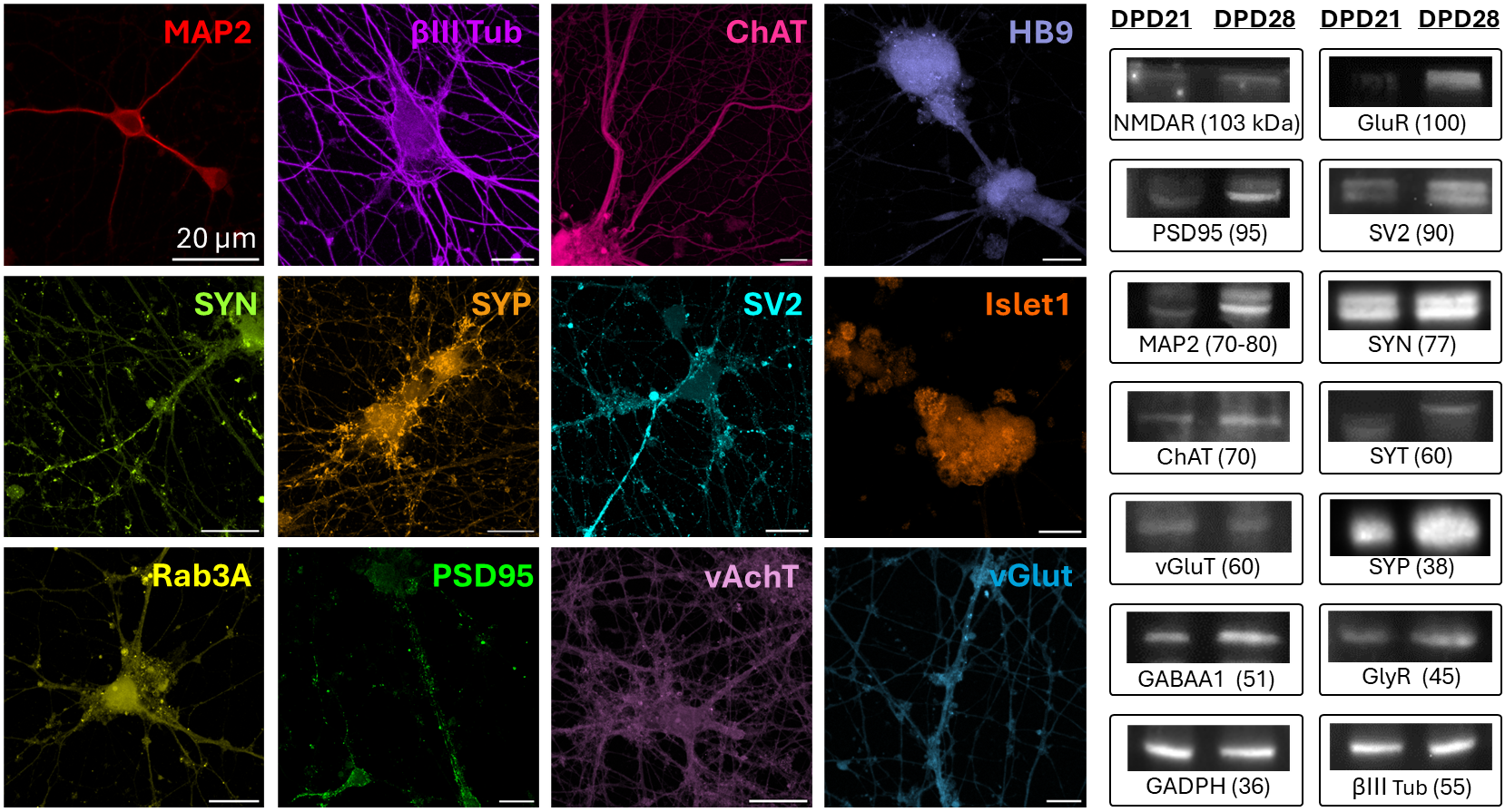
Characterization of iMNs by immunoffuorescence and immunoblotting. Immunoffuorescence staining of iMNs at DPD28 shows neuronal identity, MN specification, and synaptic maturation. Neuronal differentiation is indicated by MAP2 (red) and βIII-Tub (purple). MN identity is supported by ChAT (magenta), HBS (light purple), and Islet-1 (dark orange), which are key markers of cholinergic MNs. Synaptic development is demonstrated by the presynaptic proteins SYN (lime green), SYP (bright orange), Rab3A (yellow), and SV2 (cyan). Postsynaptic structures are labeled by PSD-S5 (bright green). Neurotransmitter identity is confirmed by the expression of vAChT (pink) and vGluT (blue), proteins responsible for acetylcholine and glutamate transport, respectively. Scale bars: 20 μm. The right panel shows corresponding western blot results, confirming increasing expression of neuronal, MN, and synaptic proteins. **Abbreviations:** MAP2: Microtubule Associated Protein 2; βIII-Tub: βIII-Tubulin; ChAT: Choline Acetyltransferase; HBS: Homeobox Protein S; Islet-1: ISL LIM Homeobox 1; SYN: Synapsin-1; SYP: Synaptophysin; Rab3A: Ras-related protein Rab-3A; SV2: Synaptic Vesicle Glycoprotein 2; PSD-S5: Postsynaptic Density Protein S5; vAChT: Vesicular Acetylcholine Transporter; vGluT: Vesicular Glutamate Transporter; NMDAR: N-methyl-D-aspartate Receptor; GluR: Glutamate Receptor; SYT: Synaptotagmin-1; GABAA1: Type A GABA (γ-aminobutyric acid) Receptor; GlyR: Glycine Receptor; GADPH: Glyceraldehyde-3-Phosphate Dehydrogenase.

**Figure 3.**
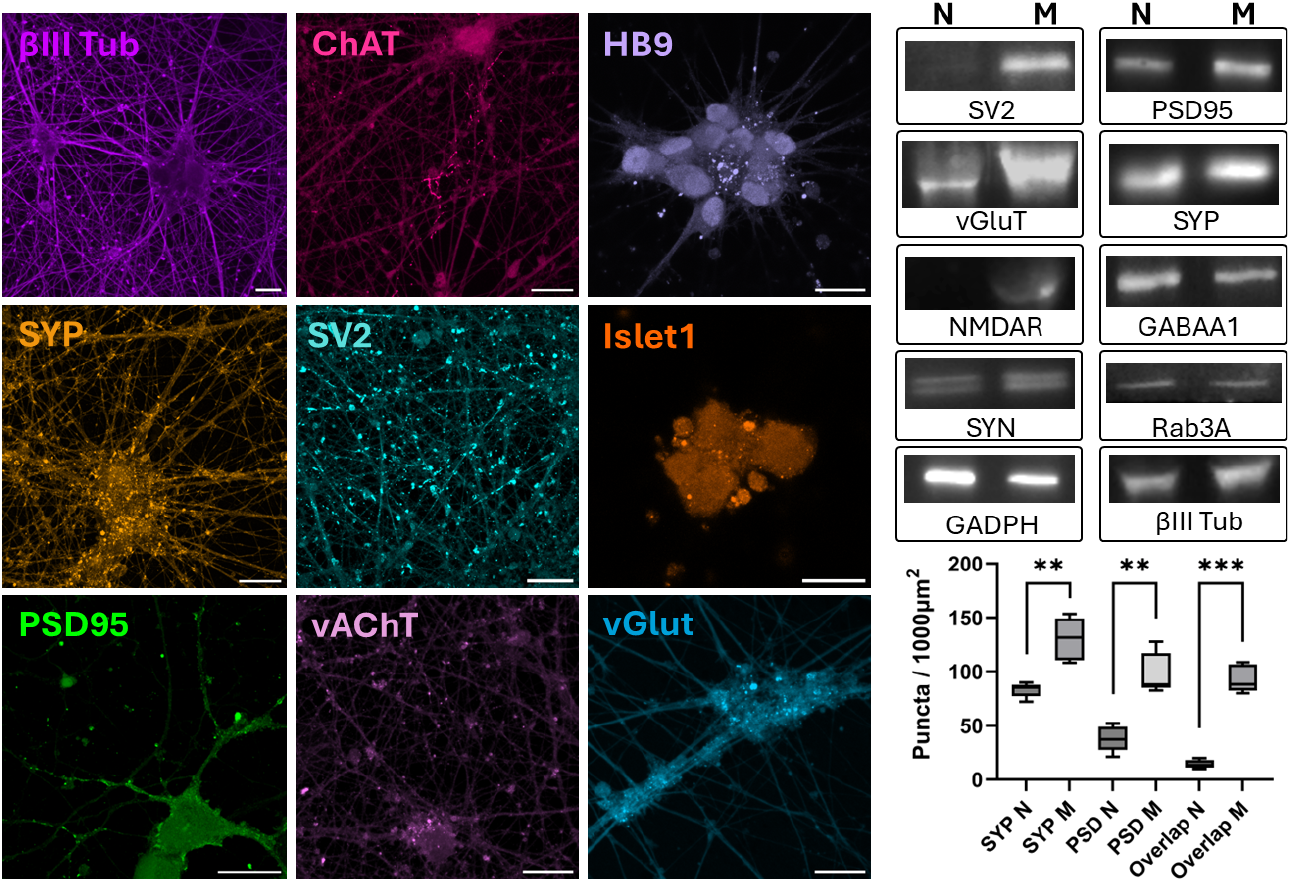
Characterization of iMNs following modulator treatment. Immunoffuorescence staining of iMNs at DPD28 shows expression of neuronal, MN, and synaptic markers following treatment with neuromodulators. Scale bars: 20 μm. See Figure 2 for a description of the labeled proteins. Western blot analysis of iMNs differentiated by the standard protocol (N) and with added neuromodulators (M) shows increased neuronal and synaptic marker expression at DPD28 for the latter (right panel, top), consistent with immunoffuorescence data. Quantification of Synaptophysin- and PSS5-positive puncta in N and M and their respective overlap (bottom right) reveals a statistically significant increase in SYP- and PSD-S5-positive puncta density as well as in overlap in modulator-treated (SYP-M) compared to untreated (SYP-N) cultures (**, p=0.00C3 (SYP); **, p=0.0038 (PSDS5); ***, p=0.0008S (Overlap), paired Student’s t-test, n=5, from 2 individual experiments). Boxplots show the median (line), interquartile range (box), and minimum/maximum values (whiskers). **Abbreviations:** see legend of Figure 2.

### Preservation of iMN Identity and Enhanced Synaptic Organization with CX516 and CDPPB

To characterize iMN identity, we stained for the MN markers HB9, Islet-1 and choline acetyl transferase (ChAT). These were abundantly expressed independently of the addition of neuromodulators, supporting the validity of our protocol (Figure 3). Qualitative comparison of synaptic markers revealed that presynaptic proteins SYP and vGluT exhibited a more punctate distribution along neurites in modulator-treated cells, while postsynaptic markers PSD-95 and GluR2 showed enhanced coalescence into discrete puncta. The latter suggests greater synaptic specialization and structural maturation in modulator-treated neurons. βIII-tubulin staining revealed increased neurite branching complexity compared to untreated controls. Treated neurons exhibited a significant increase in puncta density and pre-/postsynaptic overlap, with an average of 130.20 ± 17.56 presynaptic puncta per 1000 µm^2^ (56.6% increase, p < 0.01) and 98.50 ± 16.75 postsynaptic puncta per 1000 µm^2^ (146.5% increase). Among these, 95.5 ± 5.41% of postsynaptic puncta overlapped with presynaptic ones, and 71.85 ± 3.66% of presynaptic puncta colocalized with postsynaptic structures (n = 5 fields; Figure 3). The marked rise in postsynaptic signal (p < 0.001) was accompanied by a significant increase in synaptic overlap (p < 0.01). Together, these findings indicate that neuromodulator treatment enhances synaptic organization in iMNs without altering their motor neuron identity.

### Neuromodulators Promote an Excitatory Synaptic Phenotype

Synaptic protein expression and its temporal dynamic during maturation were assessed using western blot at DPD21 and 28 (mature state; Figure 2, right panel). All assessed proteins were detectable at DPD21 and most showed increased expression at DPD28, particularly SYN, SYP, SV2, and synaptotagmin-1 (SYT). Postsynaptic proteins, including N-methyl-D-aspartate receptor (NMDAR1), glutamate receptor (GluA2), glycine receptor (GlyR) α1, γ-aminobutyric acid type A receptor α1 (GABAA1), and PSD-95, exhibited comparatively weaker bands but also increased at DPD28. The presence of NMDAR and GluR suggests that the protocol yields iMNs enriched in glutamatergic excitatory synapses. The detection of GlyR and GABAA1 indicates the capacity to form inhibitory synapses as well, suggesting a potential excitatory–inhibitory balance necessary for network activity.

We then examined the effect of neuromodulators on synaptic protein expression at DPD28. As shown in Figure 3, top right, qualitative increases in synaptophysin, vGlut, synapsin, SV2, PSD-95, and NMDAR subunits were observed in neuromodulator-treated iMNs compared to controls. Notably, GABAA1 levels decreased after treatment, indicating a shift toward a more glutamatergic phenotype. This upregulation supports the conclusion that modulator treatment enhances synaptic protein expression, consistent with the observed increase in synaptic density.

### Progressive Acquisition of Electrophysiological Maturity Enhanced by CX516 and CDPPB

Whole-cell patch-clamp recordings were performed on iMNs at DPD7, 14, 21, 28, and 35 to assess the progression of intrinsic excitability and synaptic activity (Figure 4). From DPD14 onward, cultures were either maintained in standard differentiation medium (N) or supplemented with the glutamatergic neuromodulators CX516 and CDPPB (M).

**Figure 4.**
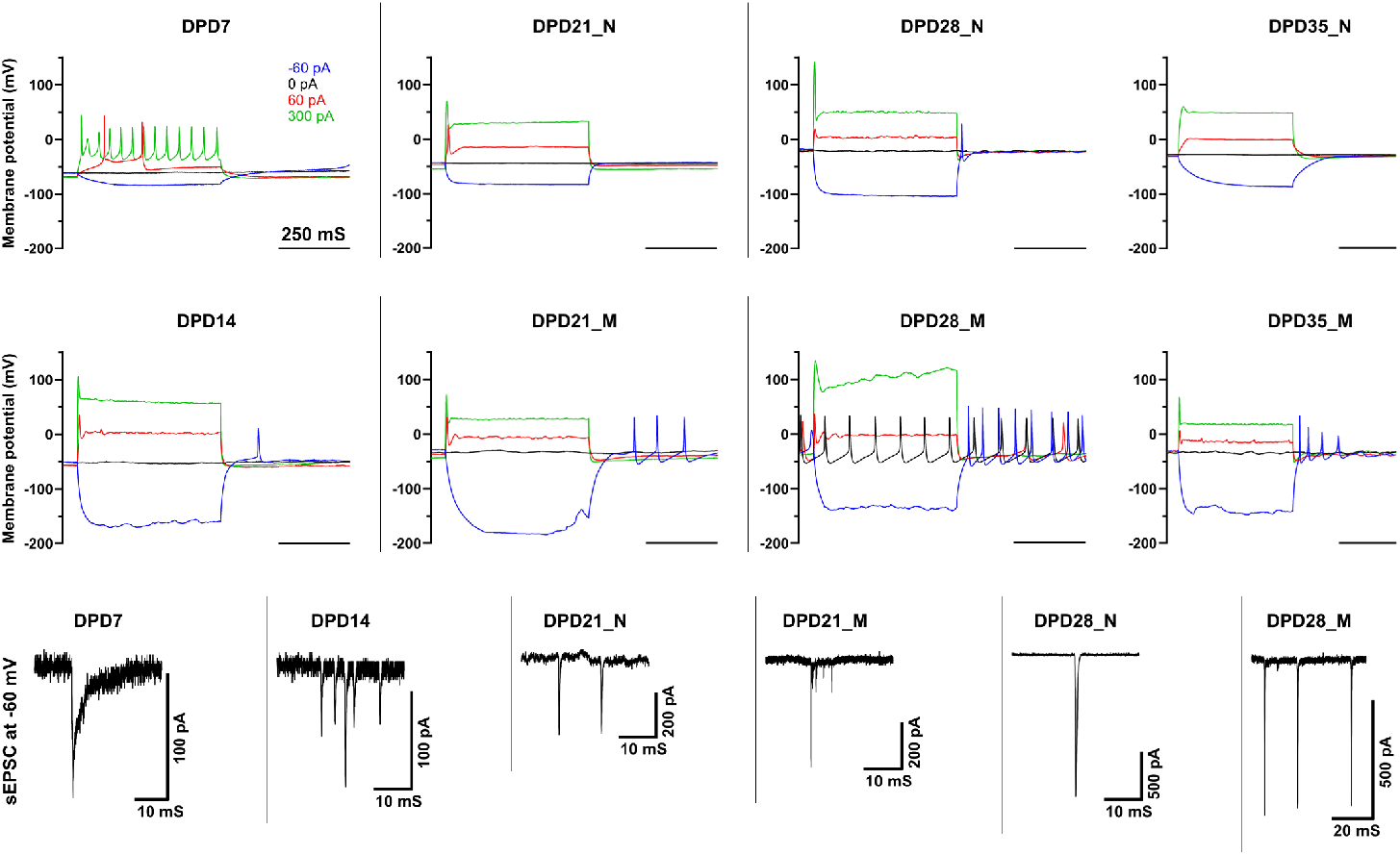
Electrophysiological maturation of iMNs over time and enhancement by CX51C and CDPPB. Whole-cell patch-clamp recordings were obtained from iMNs at DPD7, 14, 21, 28, and 35 under standard (N) and glutamatergic neuromodulator-treated (M) conditions. Representative membrane potential responses to four current injections are shown (−C0 pA, 0 pA, +C0 pA, and +300 pA), while the complete series of 20 current steps (−C0 to +320 pA in 20 pA increments) was recorded and used for quantitative analysis (Table S4). The **top** and **middle** panels illustrate the progression from early firing at DPD7 to more diverse and sustained firing patterns at later stages, with stronger repetitive activity in modulator-treated neurons at DPD28. The **bottom** panel shows representative voltage-clamp recordings of spontaneous excitatory postsynaptic currents (sEPSCs) at DPD7, 14, 21, and 28, highlighting the emergence and strengthening of spontaneous synaptic activity during differentiation (Table S5). Scale bars: 250 ms for current-clamp recordings and as indicated (10-20 ms on the x-axis; 100-500 pA on the y-axis) for voltage-clamp traces.

At DPD7, stepwise current injections already evoked multiple APs, indicating early functional expression of voltage-gated sodium and potassium channels. By DPD14-28, firing responses became more diverse, including both single and repetitive spikes, while a depolarization block appeared more frequently at stronger current injections. Quantification of AP firing patterns (Table S4) demonstrated that neuromodulator treatment significantly altered both single and repetitive AP events. At DPD21, repetitive firing during the stimulation phase showed a trend toward reduction from 6.0 ± 2.9 to 3.2 ± 0.6 APs per recording (rate ratio, RR = 0.54; 95% CI 0.27-1.08; q = 0.32), while single events were strongly increased from 36.2 ± 6.4 to 55.2 ± 8.1 APs per recording (RR = 1.52; 95% CI 1.24-1.88; q < 0.001). At DPD28, repetitive firing during the stimulation phase showed a strong trend toward increase from 1.3 ± 0.7 to 4.7 ± 2.7 APs per recording (RR = 3.50; 95% CI 1.16–10.59; q = 0.12), while single APs during the stimulation phase showed a modest decrease (42.7 → 34.3 APs per recording; RR = 0.81; 95% CI 0.62-1.04; q = 0.20).

Repetitive APs during the resting phase showed no significant change at either stage (DPD21 RR = 1.09, q = 1.00; DPD28 RR = 1.83, q = 0.30). These results indicate that CX516 and CDPPB enhance neuronal excitability in a stage- and pattern-specific manner, increasing single firing at DPD21 and promoting repetitive firing at DPD28.

In voltage-clamp mode, spontaneous excitatory postsynaptic currents (sEPSCs) were recorded to evaluate synaptic activity (Table S5). Sister cultures recorded on the same experimental dates were compared under N and M conditions. While average event frequencies showed no significant difference at DPD21 (p > 0.05), a strong increase was observed at DPD28. An incidence analysis revealed a significant enhancement in the likelihood of observing spontaneous activity in modulator-treated neurons at later stages. At DPD21, the odds of observing at least one event per run were 1.31-fold higher for M compared to N (95% CI [0.56, 3.03]; p = 0.53), whereas at DPD28, the odds increased 5.48-fold (95% CI [2.77, 10.83]; p < 0.001, FDR q < 0.001). Event frequency followed a similar pattern: no significant change at DPD21 (RR = 2.30; 95% CI [1.28, 4.14]; p = 0.005, q = 0.007), but a robust 2.51-fold increase at DPD28 (95% CI [1.72, 3.65]; p < 0.001, q < 0.001). These results indicate that neuromodulator treatment enhances both the probability and frequency of spontaneous synaptic activity, with effects becoming statistically and biologically pronounced at more mature differentiation stages.

### Calcium Imaging Confirmed Functional Responsiveness of iMNs

To confirm functional excitability through an independent assay, live calcium imaging was performed on iMNs at DPD21 using Fluo 4 AM under baseline and depolarizing conditions. Under steady-state conditions (perfusion with HEPES-ACSF), spontaneous calcium transients were detected in multiple somata and neuritic branches, reflecting ongoing network activity. Upon rapid application of high-potassium HEPES-ACSF (50 mM K^+^), a robust and near-synchronous fluorescence increase was observed across the neuronal network, indicative of depolarization-evoked calcium influx. The response was reversible, with fluorescence intensities gradually returning to baseline after stimulus removal, confirming preserved calcium homeostasis and cell viability. Representative time-lapse images illustrate these dynamics across three sequential phases: baseline, high K^+^ stimulation, and recovery (Figure 5). These observations demonstrate that the iMN cultures form a functionally mature and excitable network capable of coordinated responses to depolarizing stimuli, further corroborating the electrophysiological maturity detected in patch clamp recordings. Time-lapse videos of spontaneous and stimulus-evoked calcium activity are provided as Supplementary Movies S1a and S1b, respectively.

**Figure 5.**
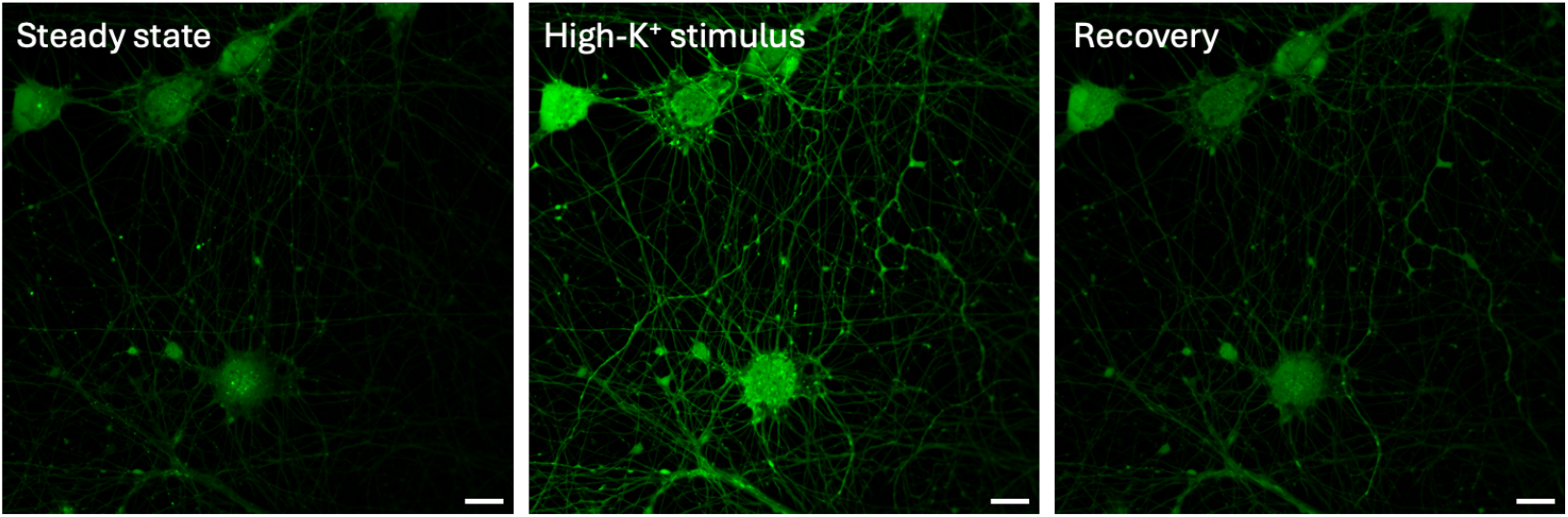
Functional calcium imaging of iMNs reveals synchronized depolarization-induced activity. Live-cell fluorescence imaging of iMNs loaded with Fluo 4 AM under baseline, depolarizing, and recovery conditions. Shown are representative frames from three key phases: (**left**) spontaneous baseline activity in HEPES ACSF (steady state), (**middle**) synchronous calcium response during perfusion with high K^+^ HEPES ACSF (50 mM K^+^), and (**right**) return to baseline following stimulus washout (recovery). Images were acquired at 2 Hz using a 40× oil immersion objective. Fluorescence intensity changes reflect intracellular calcium dynamics. Scale bar: 10 µm.

### CX516 and CDPPB Increased the Number of Recycling SVs Without Altering Release Kinetics

AM4-64 live imaging revealed evocable SV recycling in iMNs. High-potassium depolarization in the presence of AM4-64 produced fluorescent puncta along neurites, reflecting dye uptake during endocytosis. Subsequent depolarization in dye-free high-potassium solution led to destaining, consistent with exocytosis of recycling SVs (Figure 6, top left graph). By DPD28, iMNs displayed robust staining/destaining profiles characteristic of functional synapses. CX516 and CDPPB treatment significantly increased the number of AM4-64-labeled puncta undergoing destaining compared to controls, indicating a higher number of active presynaptic sites (Figure 6, bottom right panel, ∼2.5× increase, P<0.0001). This contributes to enhanced synaptogenesis and greater availability of recycling vesicle pools. However, the fluorescence decay kinetics remained unchanged, with overlapping curves and similar release rates (κ), indicating that synaptic modulators did not alter the speed or extent of vesicle turnover, while effectively increasing synaptic density (Figure S2).

**Figure 6.**
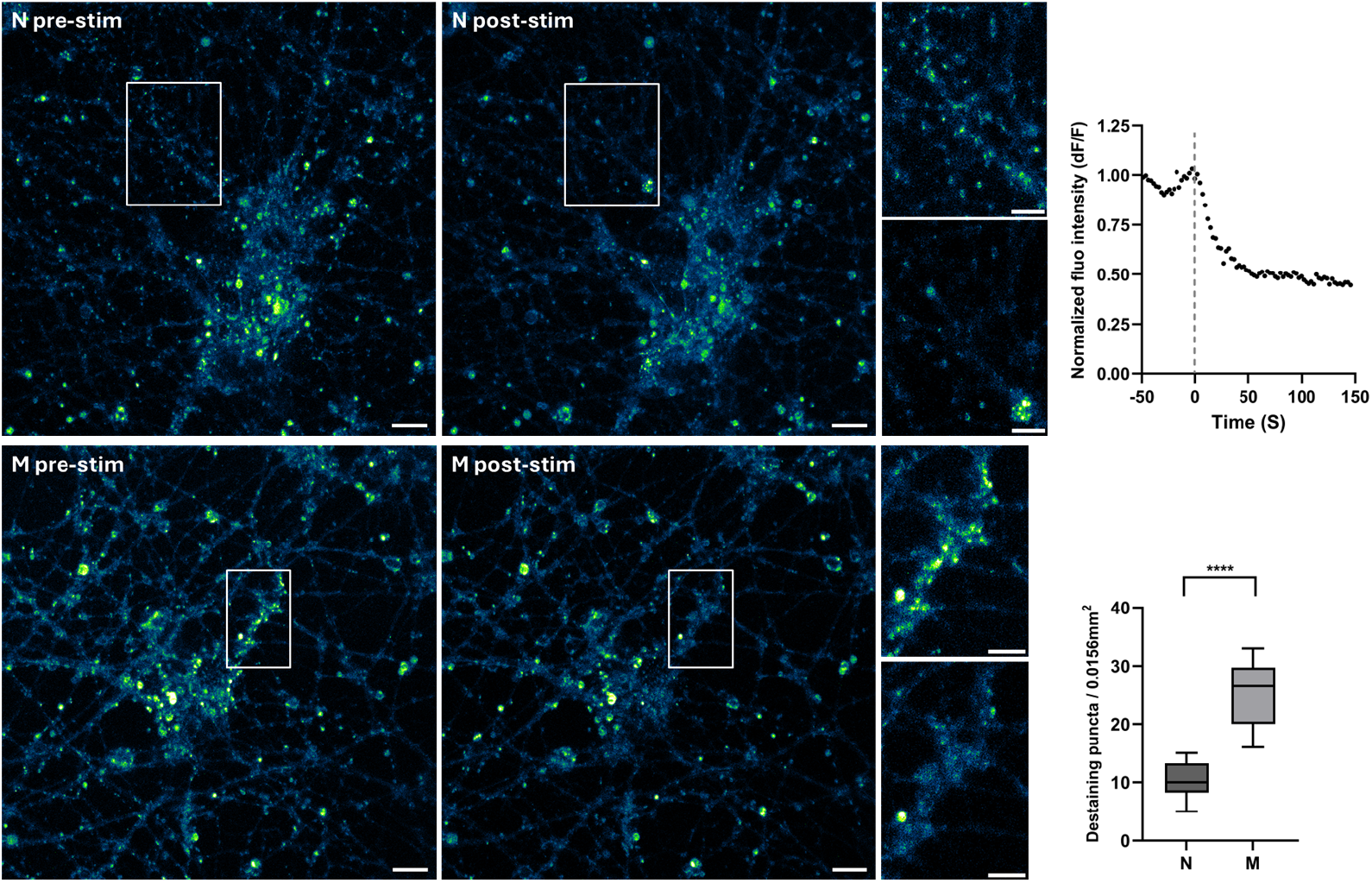
Functional live staining assay shows significant increase in destaining puncta upon neuromodulator treatment. iMNs are differentiated by the standard protocol (N; **top row**) and with neuromodulators added (M; **bottom row**) both pre- (N/M pre-stim) and post-high-KCl stimulation (N/M post-stim) with respective insets. The left panel displays the stained state prior to the stimulation of evoking destaining, with numerous bright puncta representing dye-loaded, active presynaptic terminals. Following high-potassium stimulation (middle panel), a noticeable reduction in AM4-64-positive puncta is observed, indicating dye release and thus SV exocytosis. Insets (right montages) show magnified regions highlighting the decrease in puncta intensity post-stimulation. Scale bars for full images and insets, 10 and 3 µm, respectively. **Top right panel**: Fluorescence decay curve of high-KCl stimulation in iMNs (N), mean intensity of all puncta of one example field-of-view, indicating robust SV exocytosis. **Bottom right**: Quantification of destaining puncta following KCl-induced exocytosis in normal (N) and modulator-treated (M) iMNs. Boxplot analysis of AM4-64 destaining puncta per 0.0156 mm^2^ shows a significant increase in active exocytosis sites in modulator-treated neurons compared to untreated controls. The median number of destaining puncta is markedly elevated in the modulator group, indicating an increase in the number of functional synapses. Statistical analysis reveals a highly significant difference between the two groups (p < 0.0001, paired Student’s t-test, n=8, from 3 individual experiments).

### Ultrastructural Features of iMNs and Effects of Glutamatergic Modulation

Conventional EM was used to examine ultrastructural development of iMNs over a 35-day differentiation period. Up to day 28, neuronal cell bodies maintained intact morphology, including well-defined somata and clear nuclear profiles (Figure S3, left micrograph). By day 35, however, signs of cellular stress, such as cell detachment and fragmentation, became apparent under brightfield observation in the culture dish. These visual cues were associated with increased difficulty during subsequent sample preparation. These findings are consistent with previous reports of stress responses in long-term iPSC-derived neuronal culture^37,38^.

At the synaptic level, presynaptic boutons were readily identifiable and contained dense clusters of SVs together with defined active zones. Adjacent postsynaptic densities were present, forming morphologically mature synaptic contacts. Large dense-core vesicles (60 to 100 nm in diameter) were detected in varicosities (Figure S3, right micrograph), consistent with storage of neuropeptides involved in synaptogenesis and synaptic plasticity^39–41^. Elongated mitochondria were frequently observed in neuronal processes, consistent with the high metabolic demands of MNs (Figure 7, bottom middle micrograph).

**Figure 7.**
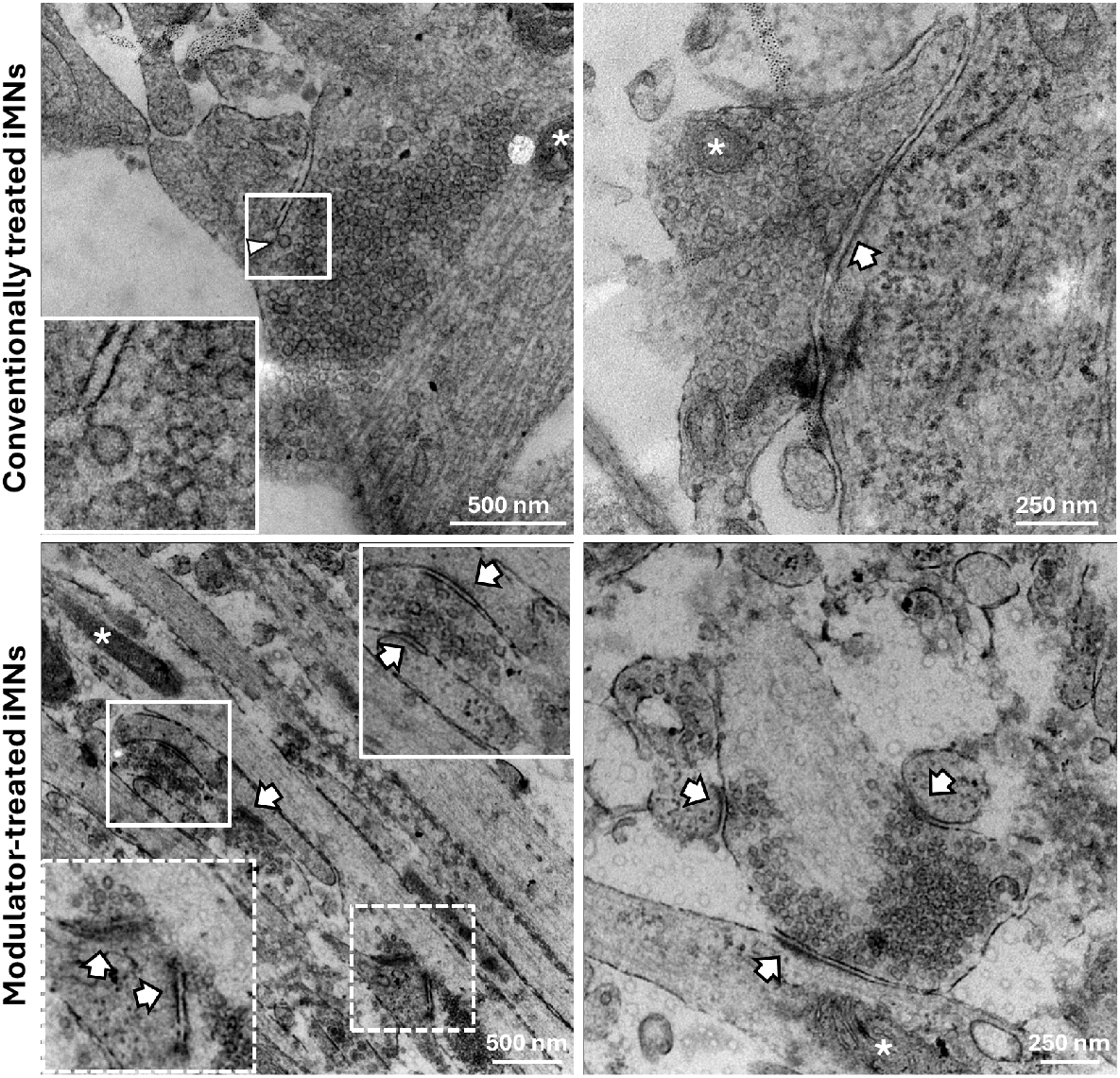
Transmission electron microscopy (EM) of presynaptic terminals in iMNs. Ultrathin sections show synaptic boutons densely packed with synaptic vesicles (SVs), including vesicles adjacent to the presynaptic membrane, indicative of docking at active zones. Each panel highlights a distinct synapse, demonstrating variability in bouton morphology and vesicle organization. Synaptic clefts are visible in several micrographs, separating pre- and postsynaptic membranes. **Top panel:** A recycling SV is marked by a white arrowhead. Mitochondria (asterisk) are occasionally observed near vesicle clusters. The presence of round vesicles, defined active zones, and synaptic clefts is consistent with functional excitatory synapses^43^. **Bottom panel:** EM images of modulator-treated iMNs reveal enhanced active zone formation. Presynaptic terminals display densely packed SVs and extended, electron-dense synaptic clefts (arrows located at postsynaptic zone), more prominent than untreated iMNs. Insets show higher magnification views of active zones aligned with postsynaptic densities. The right panel illustrates bouton regions with vesicle clusters tightly positioned near thickened presynaptic membranes, consistent with mature active zone architecture. Scale bars: 500 nm (left and middle), 250 nm (right).

In cultures treated with CX516 and CDPPB, synaptic boutons were more frequently observed and appeared structurally more complex than in control iMNs (Figure 3 and 7). Notably, some boutons displayed multiple active zones, accompanied by a higher number of readily releasable SVs tightly arranged at the presynaptic membrane, features associated with increased synaptic connectivity^42^ (Figure S4 left panel). These findings indicate that glutamatergic modulators promote the formation of more abundant and structurally more elaborate boutons in iMNs, in agreement with the functional assays described above.

While conventional EM confirmed mature synaptic structures in iMNs, this approach is limited by chemical fixation and heavy-metal staining, which can obscure finer ultrastructural details. To capture the native organization of synapses and subcellular compartments, we turned to cryo-ET. For this, iMNs were differentiated directly on EM grids and vitrified without chemical fixation, enabling three-dimensional visualization of their architecture in a near-native state.

A key challenge was obtaining samples thin enough for tomography without resorting to cryo-FIB milling. Prolonged differentiation led to dense neuronal networks that retained liquid and frequently produced ice that was too thick for imaging. By implementing modified blotting strategies, we reproducibly improved sample preparation outcomes, with on average ∼50% of the 22 prepared grids providing sufficient regions for cryo-ET data collection. (Figure S5; see Materials and Methods). This optimization enabled high-quality cryo-ET of presynaptic terminals and neuronal processes, preserving ultrastructural details.

High-resolution cryo-ET revealed the organization of synaptic and subcellular compartments (Figure 8). Synaptic boutons were found densely filled with uniformly sized SVs. Postsynaptic densities and synaptic clefts, though less distinctly contrasted than in conventional EM, were nevertheless consistent with functional synaptic contacts. Small synapses were also observed along axons, characterized by SV tethering to the plasma membrane, potentially representing *en passant* synapses or developing contact sites indicative of MN maturation and synaptic integration (Figure 8, middle panel and figure S4 right panel). This distribution is consistent with the distinct connectivity of MNs, which can form both terminal and *en passant* synapses^44–46^.

**Figure 8.**
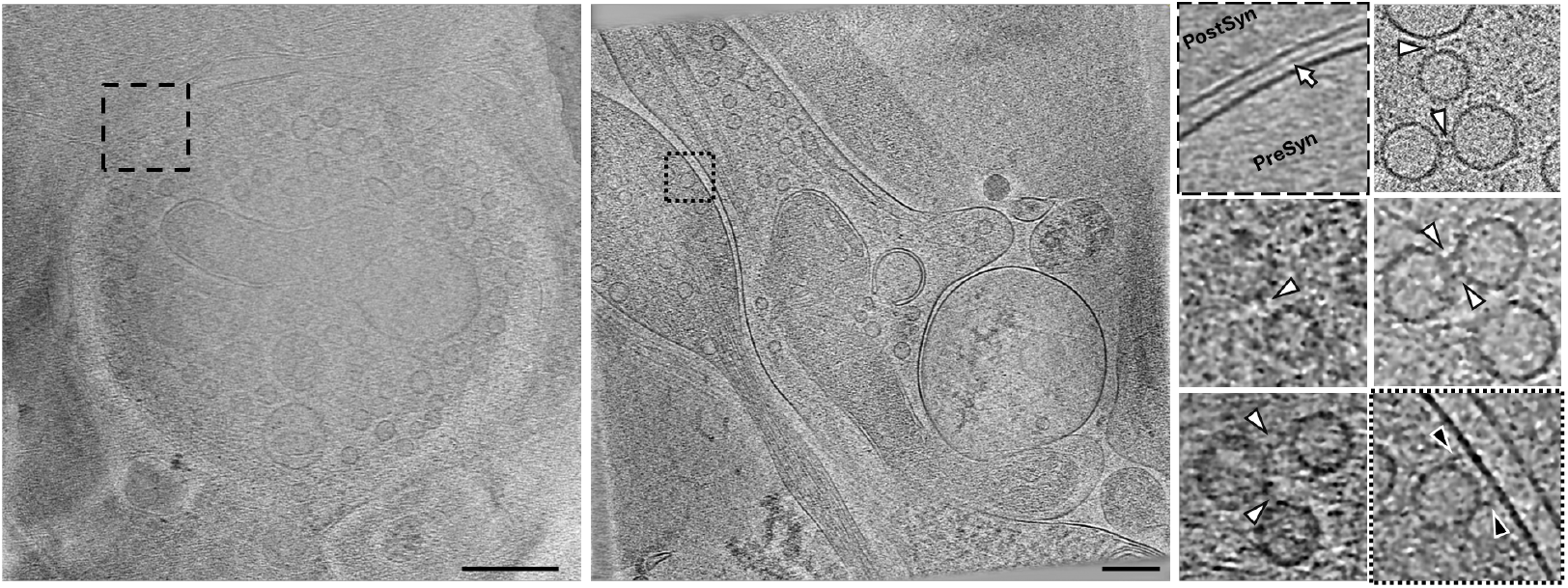
Cryo-ET of a presynaptic bouton in iMNs. Cryo-ET reveals the three-dimensional ultrastructure of presynaptic terminals enriched with SVs. **Left panel:** A tomographic slice shows a bouton with abundant SVs densely clustered near the interface with the postsynaptic compartment. A centrally located mitochondrion is visible. **Middle panel:** A varicosity-like presynaptic structure displays a dense SV cluster, with vesicles tethered to the plasma membrane. Scale bars: 200 nm. **Right panel** (montage of magnified subregions from the same datasets)**: Top left:** The synaptic cleft (∼20 nm) separates the presynaptic and postsynaptic membranes (PreSyn and PostSyn). The white arrow highlights electron-dense material spanning the cleft, potentially corresponding to adhesion molecules or structural components of the trans-synaptic complex. In the other subregions, white-filled arrowheads indicate inter-vesicular connectors, likely representing proteinaceous linkers. Black-filled arrowheads denote tethers anchoring vesicles to the plasma membrane, marking the docking stage and possibly priming for fusion.

Cytoskeletal organization was well resolved in axonal and dendritic processes (Figure S6). Parallel bundles of microtubules and neurofilaments were visible. In addition to individual actin filaments, bundled actin structures were also observed. Elliptical vesicles were frequently observed along microtubules, suggesting active transport. Multivesicular bodies decorated with membrane proteins were identified. Lysosomes and autolysosomes were present at various stages of degradation, and occasionally extracellular lysosome-like features suggested release by either cell rupture^47,48^. Mitochondria with distinct cristae were abundant in processes, while smooth endoplasmic reticulum was distributed throughout the extensions.

Tomograms further resolved SV spatial organization within presynaptic boutons. They exhibited densities tethering them to the plasma membrane, and they were attached to each other by connectors (Figure 8) as observed in rodent neurons^8,49,50^.

To relate ultrastructure to function, we used cationized ferritin as an electron-dense tracer, which binds electrostatistically to the plasma membrane. Following high-potassium stimulation, ferritin was internalized into endosomes of 50-200 nm, consistent with synaptic activity-dependent endocytosis (Figure S7)^51^.

Taken together, our electrophysiological, imaging, and ultrastructural analyses demonstrate that iMNs progressively acquire hallmark features of functional MNs, including excitability, synaptic connectivity, and SV recycling. Modulation of glutamatergic receptors further enhanced synaptic density and bouton complexity without altering SV release kinetics. Conventional EM confirmed the presence of presynaptic terminals, synaptic clefts, postsynaptic specializations, and dense-core vesicles, while cryo-ET provided high-resolution views of synaptic architecture and cytoskeletal organization in a near-native state. Functional assays using AM4-64 dye and cationized ferritin revealed activity-dependent endocytosis and evocable exocytosis, underscoring neuronal maturity and competence. These findings validate our protocol for generating iMNs as a reliable model to study human synaptic physiology and to explore mechanisms relevant to neurodegenerative disease.

## Discussion

Our study establishes a reproducible and efficient method for differentiating human iMNs, providing a platform to investigate synaptic mechanisms relevant to MN physiology and disease. We validate their synaptogenic capacity using a multimodal approach combining immunostaining, electrophysiology, live-cell imaging, and EM, demonstrating that iMNs differentiated by this method serve as a suitable in vitro model for studying synaptic function.

Initial analyses showed that, although cellular interconnections were evident, the number of synaptic contacts was lower than expected, as shown in functional, immunofluorescence, and ultrastructural assays. This limitation reflects a common challenge in neuronal culture systems: maintaining long-term cell adhesion and promoting robust synaptogenesis. This issue is more pronounced in human iPSC-derived neurons, which often require extended maturation times as compared to primary neuronal cultures.

To address this, we applied a combination of two glutamatergic modulators, the AMPA receptor potentiator CX516 and the mGluR5 modulator CDPPB. Both have been reported in other systems to enhance plasticity and rescue synaptic or behavioral deficits^29,30,52^. In our experiments, their combined application promoted synaptogenesis in iMNs, as shown by increased expression of synaptic proteins, higher synaptic densities, and more complex presynaptic boutons. Importantly, these effects were achieved without altering MN identity, underscoring the feasibility of using neuromodulation to improve functional maturation of human iMNs in vitro. Nonetheless, variability in synapse density across samples highlights the need for further optimization of culture conditions, modulator combinations, and dosages.

An important limitation of our study is inherent to the use of neuromodulators to promote synaptogenesis. On the one hand, such compounds accelerate or augment synapse formation through pathways that may not reflect the physiological developmental trajectory of MNs. This could reduce interpretability in disease-modeling contexts, where altered development may itself be part of the pathology. On the other hand, neuromodulator treatment can correct for intrinsic limitations of iMN cultures, such as slow or incomplete synaptogenesis, thereby enabling ultrastructural analysis of synapses that would otherwise be too sparse. Notably, our data show that SV recycling dynamics are comparable between treated and untreated cultures, suggesting that presynaptic function can be meaningfully interpreted in disease models, even while the absolute number of synapses has been pharmacologically increased.

In a broader context, our findings provide a framework for future research into synaptic mechanisms in MN diseases. The successful application of cryo-ET in iMNs demonstrates the feasibility of capturing near-native ultrastructure of cultured human synapses, complementing functional assays and paving the way for studies of synaptopathies at molecular resolution.

Finally, we also highlight technical challenges of applying cryo-ET to mature neurons cultured directly on grids. While cryo-FIB milling enables thinning of thick regions, it remains difficult to apply reliably to synaptic regions due to their small size and the limited sensitivity of integrated fluorescence systems for targeting. In addition, differentiating neuronal cultures directly on grids often generates samples that are too thick for tomography, as prolonged culture leads to dense neurite bundles. These thick regions impair blotting and vitrification, frequently yielding unusable ice, and remain problematic even when FIB milling is attempted. To address these issues, we optimized grid preparation and blotting conditions, enabling long-term adhesion of iMNs and improved vitrification without the need for FIB milling. This strategy reproducibly yielded thin, well-preserved neuronal regions suitable for high-quality cryo-ET, thereby allowing direct ultrastructural analysis of synapses in human iMNs.

In conclusion, we show that human iMNs differentiated on EM grids, combined with glutamatergic modulation, develop sufficient synaptic density and structural complexity to permit detailed functional and ultrastructural studies. This platform provides a robust basis for investigating human synaptic physiology and its alterations in neurodegenerative disease models.

## Materials G Methods

### iMN Culture

The iPSC lines were obtained from WiCell and were kindly gifted by Saxena Smitas’ laboratory (two healthy lines)^53^. The iPSC lines were cultured on Geltrex™ (Thermo Fisher Scientific) coated dishes at 37°C with 5% CO_2_ and maintained in mTeSR™ Plus medium (StemCell technologies). The medium was supplemented with 10 μM Y-27621 dihydrochloride (RI (rock inhibitor)) (StemCell technologies) when cells were split. Medium was exchanged the day following splitting without RI, maintaining cells by exchanging 2mL mTeSR™ Plus every second day until 80% confluence was reached. MN differentiation was adapted from previously established protocols, with modifications introduced to optimize the procedure for our experimental conditions^53–55^. To initiate embryoid body (EB) formation, iPSCs were dissociated into small clusters using EZ-LiFT™ reagent (Sigma), and 3×106 cells were seeded onto 60 mm dishes with N2B27 differentiation medium (Neurobasal Plus: Advanced DMEM/F12 (1:1), 1x Pen/Strep (Gibco), 0.1 mM 2-mercaptoethanol (Gibco), supplemented with 1x N2 supplement (Gibco), 1x B27 plus supplement (Gibco), 10 μM RI, 10 ng/mL basic fibroblast growth factor (Sigma), 10 μM L-ascorbic acid (LAA; Sigma), 10 μM SB431542 (SB; Sigma), 3 μM CHIR99021 (CHIR; Sigma), and 0.1 μM LDN193189 (LDN; Sigma). On day 2, patterning was induced by adding 10 μM L-ascorbic acid, 10 μM SB (Sigma), 3 μM CHIR, and 0.1 μM LDN193189 until day 7, as well as 500 nM smoothened agonist (SAG; Sigma) and 100 nM all-trans retinoic acid (RA; Sigma) until day 16. 10 ng/mL brain-derived neurotrophic factor (BDNF; Sigma), 10 μM DAPT (Sigma), and 10 ng/mL glial-derived neurotrophic factor (GDNF; Sigma) were added from day 7, 9, and 14, respectively. After 16 days, EBs were dissociated using trypsin and triturated with HD-FBS, both supplemented with DNase. Triturated cells were seeded on poly-L-ornithine (0.01%) / laminin (1/333)-coated dishes and maintained in MN feeding medium consisting of Neurobasal Plus, 1x non-essential amino acids (NEAA, Thermo Fisher Scientific), 1x Pen/Strep, and 0.1 mM 2-mercaptoethanol supplemented with 1x N2, 1x B27, 10 μM LAA, 100 nM RA, 10 ng/mL BDNF, 10 ng/mL GDNF, 10 ng/mL ciliary neurotrophic factor (CNTF; Sigma), and 10 ng/mL insulin-like growth factor (IGF-1; Sigma). To enhance synapse maturation, the glutamatergic modulators CX516 and CDPPB were added 14 days after MN differentiation onset (DPD14) and tested at three concentrations (100 nM, 1 μM, 10 μM). The lowest concentration (100 nM) was chosen due to neurotoxic effects at higher doses (see Supplementary Figure S8). Medium was exchanged every 3-4 days, and cells cultured for up to 28 days and processed for downstream experiments. More details are available in the Supplementary Materials.

Maintaining long-term cell adhesion presents a significant technical challenge in post-mitotic neuronal cultures, particularly for human neurons^37^. Unlike primary animal neurons, human neurons require extended maturation periods to establish functional synaptic networks in vitro^56^. This prolonged timeline increases the risk of detachment and aggregation, posing ongoing difficulties for which effective solutions remain limited. Aggregation of somata and neurites can be reduced by complete EB disruption, requiring adequate amounts of trypsin and DNase, and by seeding an appropriate cell density, while preventing spontaneous differentiation. The latter can be suppressed by adding 1–2 μM 5-Fluorodeoxyuridine (FdU) (Sigma Aldrich) or arabinocytosine (AraC) (Sigma Aldrich)^57^. Depending on cell seeding density, neurites may form a carpet-like mat, which is more prone to film-like detachment. Thusa lower seeding density of 500,000 cells per well in a 6-well plate is advised. Cell detachment during prolonged culture can be reduced by supplementing the medium with laminin at a 1:1000 dilution every other medium exchange. In cell lines with low survival after plating, we advise increasing the seeding density by 20-30%.

### Immunofluorescence Microscopy

For immunostaining experiments, cells were fixed using a mixture of 3% paraformaldehyde (PFA) and 0.5% glutaraldehyde (GA) in HEPES buffer for one hour at room temperature. This was followed by three washes in phosphate-buffered saline (PBS). The combination of PFA and GA provides high-quality fixation, while allowing the same samples to be used for conventional EM. Cells were permeabilized with 0.3% Triton X-100 for 15 min and washed again with PBS. 3% SureBlock (LubioScience) in PBS was used for blocking for 1h at room temperature. Cells were labeled with primary and subsequently secondary antibodies (see Table S2) in 3% SureBlock for 1h at RT, washing in between with PBS. The coverslips were dried overnight at 4°C and then mounted on glass slides using ProLong Diamond Antifade Mountant (Thermo Fisher Scientific) and left to harden at room temperature for 24h. The mounted coverslips were stored at 4°C until they were imaged using a confocal Zeiss LSM 880. For statistical analyses, images were analyzed as maximum-projected Z-stacks. Puncta were counted using a custom-made macro in Fiji^58^.

### Western Blot

For immunoblots, neurons were cultured as mentioned in a 60 mm dish at a seeding density of 1.5x 10^6^ per dish. Cells were washed and lyzed using RIPA buffer (50 mM Tris-HCl, pH 7.4, 150 mM NaCl, 1% Triton X-100, 0.5% Na-deoxycholate, 0.1% SDS) with 1:100 protease inhibitor cocktail (Sigma Aldrich). Cells were incubated for 30 min on ice, sonicated, and further incubated on ice for 10 min. The lysed cells were centrifuged at 14,500x g for 20 min, and the supernatant containing the proteins of interest was collected and frozen (−80°C) until use^59^. The protein content was determined using a BCA assay (Thermo Fisher Scientific). Equal amounts (5μg) of total protein were separated on 10% SDS-PAGE gels and transferred onto nitrocellulose membranes (Millipore). The membrane washed with PBS containing 0.1% Tween 20 (PBST) for 10 min and then incubated with Intercept blocking buffer (LiCOR) in PBST for 1h. Incubation with primary antibodies at a 1:5000 dilution in blocking buffer was performed overnight at 4°C, washed 3x with PBST and then further incubated with HRP-conjugated secondary antibodies for 1h. After a final wash, the membranes were incubated with a luminol-based detection reagent, and the signal was captured using the Fusion Fx system (Vilber Lourmat).

### Whole-Cell Patch-Clamp Electrophysiology and Analysis

iMNs cultured on 12 mm glass coverslips were transferred to a recording chamber perfused with room-temperature, oxygenated, carbonated artificial cerebrospinal fluid (ACSF; in mM: 125 NaCl, 2.5 KCl, 25 NaHCO_3_, 1.25 NaH_2_PO4, 1 MgCl_2_, 2 CaCl_2_, 25 glucose). Whole-cell patch-clamp recordings were conducted using heat-pulled borosilicate glass pipettes (4-8 MΩ) filled with intracellular solution containing (in mM): 130 K-gluconate, 5 KCl, 10 Na-phosphocreatine, 4 Mg-ATP, 0.3 Na-GTP, and 10 HEPES (pH 7.2). MNs were visualized using an infrared CCD camera mounted on a Leica DM LFSA microscope.

For current-clamp recordings, square current pulses of 500 ms duration were injected in 20 pA increments from −60 pA to +320 pA to elicit action potentials (APs). The complete series of 20 current steps was used for quantification. Firing responses were categorized as repetitive or single APs during the stimulation phase (SP) and as repetitive or single APs during the resting phase (RP).

For each coverslip, the total number of APs across all current steps and patched cells was counted. Event rates were expressed as mean APs per recording and reported as mean ± SEM (Table S4). To evaluate modulation by CX516 + CDPPB, simple per-step event-rate comparisons between neuromodulator-treated (M) and untreated (N) cultures were performed using a Poisson-type rate ratio model with exposure defined as the total number of injected current steps (20 steps × number of coverslips per condition). The rate ratio was computed as RR = (y_M / E_M) / (y_N / E_N), where y is the total event count and E the exposure. Approximate 95% confidence intervals were calculated as exp[ln(RR) ± 1.96 × √(1/y_M + 1/y_N)], with Haldane-Anscombe correction (+0.5) applied to both counts if any y = 0. Two-sided p-values were derived from the Wald statistic of the log-rate ratio. p-values were adjusted using Benjamini-Hochberg FDR within each developmental stage (four comparisons: repetitive APs during the stimulation phase (rSP), single APs during the stimulation phase (sSP), repetitive APs during the resting phase (rRP), single APs during the resting phase (sRP)). This approach provided a robust measure of neuromodulator-induced changes in firing frequency across differentiation stages.

For voltage-clamp recordings, spontaneous excitatory postsynaptic currents (sEPSCs) were recorded at a holding potential of −60 mV in oxygenated aCSF. Event detection was performed using a 5 pA threshold in Igor Pro (WaveMetrics). For consistency, prominent sEPSC peaks exceeding 100 pA were manually selected for each recording to ensure reliable amplitude-based identification. For each cell, mean event frequency and positivity (fraction of active 1 s epochs containing at least one event) were calculated.

To assess the incidence of spontaneous activity, the probability of observing at least one event per run was modeled using a binomial generalized linear model (GLM) with a logit link. Each cell contributed *k*_*i*_ active runs out of *n*_*i*_ total runs (*k*_*i*_ *∼ Binomial(n*_*i*_, *p*_*i*_)). Condition effects were estimated by maximum likelihood, controlling for paired dates as a categorical factor, and odds ratios (OR) were derived as e^β with 95% confidence intervals. Significance was tested using two-sided Wald tests and corrected for multiple comparisons using the Benjamini-Hochberg false discovery rate (FDR) method.

For event frequency, a Poisson GLM with log link was used for spike counts, with offset = log(n_*i*_). Rate ratios (RR) were derived as e^β with 95% CI and Wald p-values, FDR-corrected.

Comparisons between untreated (N) and neuromodulator-treated (M) cultures were performed between paired sister cultures recorded on the same experimental dates (Table S5). Graphs and statistical analyses were performed using Igor Pro, R, and GraphPad Prism.

### Calcium Imaging

Calcium imaging experiments were performed using standard HEPES-buffered artificial cerebrospinal fluid (HEPES-ACSF) containing (in mM): 140 NaCl, 5 KCl, 2 CaCl_2_, 1 MgCl_2_, 10 HEPES, and 10 D-glucose. The pH was adjusted to 7.4 with NaOH, and osmolarity was maintained at approximately 300 mOsm. For depolarization experiments, a high-potassium variant (high-K^+^ HEPES-ACSF) was prepared by replacing NaCl with 50 mM KCl and reducing NaCl to 95 mM to maintain osmotic balance. All solutions were prepared with ultrapure water and filtered through a 0.22 µm membrane. Standard HEPES-ACSF was stored at 4 °C for up to one week, while high-K^+^ HEPES-ACSF was freshly prepared before each experiment. The dye loading solution was prepared immediately before use by dissolving the calcium-sensitive AM-ester dye Fluo-4 AM (Biotium) to a final concentration of 5 µM in HEPES-ACSF supplemented with 0.02% (w/v) Pluronic® F-127 (Sigma Aldrich) (from a 20% stock in DMSO). iMNs cultured on glass-bottom dishes were rinsed once with HEPES-ACSF and incubated with the dye loading solution for 30 min at 37 °C in the dark. Following incubation, cells were washed three times with dye-free HEPES-ACSF and allowed to de-esterify for 15–20 min at room temperature before imaging.

Live calcium imaging was performed using a Zeiss LSM 880 confocal microscope equipped with a 40× oil immersion objective and environmental control maintained at 32–35 °C. Excitation was achieved at 488 nm, and emission was collected between 500–550 nm. Laser intensity and detector gain were minimized to prevent photobleaching and saturation. The pinhole was set to 2 Airy units, and images were acquired at 2 Hz. The recording sequence comprised three phases: 1) Baseline (spontaneous activity): 2 min in standard HEPES-ACSF, 2) Depolarization: rapid perfusion with high K^+^ HEPES-ACSF (50 mM K^+^) for 2 min, 3) Recovery: washout with standard HEPES-ACSF and continued recording for 3 min.

### AM4-64 Live-Cell Staining and Destaining

To evaluate synaptic activity and vesicle recycling, live-cell staining was performed using the fluorescent membrane dye AM4-64 (Biotium, VWR International). The staining protocol was adapted from Iwabuchi et al. and Ratnayaka et al^60,61^. iMNs were cultured on µ-dishes (ibidi) and imaged in a temperature-controlled perfusion chamber at 33-35 °C in Tyrode’s solution (in mM: 124 NaCl, 5 KCl, 2 CaCl_2_, 1 MgCl_2_, 30 glucose, and 25 HEPES, pH 7.4). SV recycling was triggered by depolarization with high-potassium Tyrode’s solution (70 mM KCl, 59 mM NaCl) containing AM4-64, which was taken up into recycling SVs via endocytosis. Non-internalized dye was quenched with SCAS (Biotium) in low-calcium Tyrode’s solution. Time-lapse imaging was conducted at 0.5 Hz during a second high-potassium stimulation (without dye) to record distaining of previously labeled SVs undergoing exocytosis. Z-stacks were acquired pre- and post-stimulation to control z-axis movement-related signal changes. Fluorescence decay curves were generated from time-lapse data at active sites and fitted with an exponential decay model.

### Conventional EM

After removal of previous media, cells were submerged in fixative (2.5% glutaraldehyde (Agar Scientific) in 0.15M HEPES (Fluka), 670 mOsm, pH 7.35) at 4°C for at least 24h. They were then washed with 0.15 M HEPES three times for 5 min, postfixed with 1% OsO4 (EMS) in 0.1M Na-cacodylate-buffer at 4°C for 1h. Thereafter, cells were washed in 0.1 M Na-cacodylate-buffer three times for 5 min and dehydrated in 70, 80, and 96% ethanol (Alcosuisse) for 15 min each at room temperature. Subsequently, cells were immersed in 100% ethanol (Merck) three times for 10 min, and then submerged with ethanol-Epon (1:1) overnight at room temperature. The next day, cells were embedded in Epon (Fluka) and left to harden at 60°C for 5 days.

Sections were prepared using a UC6 ultramicrotome (Leica), beginning with 1 μm semithin sections for light microscopy, stained with 0.5% toluidine blue O (Merck). Ultrathin sections (70-80 nm) were then produced for EM. These sections were mounted on copper grids, including 200-mesh square, single-slot, and 100-mesh hexagonal grids. They were stained with uranyless (EMS) and lead citrate (Leica) in an EMstain (Leica). Imaging was conducted at magnifications between 18,000x and 30,000x with a Tecnai Spirit G2 electron microscope (FEI) operating at an acceleration voltage of 80 kV, and images were captured using an Olympus-SIS Veleta CCD camera.

### Cryo-ET

Cryo-ET experiments were performed on gold EM grids using standard cell culture methods with specific modifications. Table S3 presents the grid types tested, the challenges encountered, the adaptations implemented, and their overall suitability for cryo-ET sample preparation. Key challenges included achieving proper cell adherence and ensuring adequate thinness of the grids for high-quality imaging. Grids were first coated with poly-L-ornithine (PLO) and Laminin, both diluted in Neurobasal medium (0.01% and 1/333, respectively), then placed in culture dishes and incubated at 37 °C for 1 hour prior to cell seeding. This preparation improved grid handling and ensured better cell attachment during the seeding process. Prior to cryo-fixation, 4 µL of 10 nm uncoated gold nanoparticles (GoldSol; Aurion) were applied as fiducial markers. Vitrification using standard blotting conditions yielded excessively thick ice, in particular in long-differentiated cultures. Grids were first blotted from the back, cell-free side for 6-8 seconds (Figure S5). A supplementary front-blotting step (2-3 seconds) was then performed using the torn edge of a filter paper held perpendicular to the grid surface near the tweezers, which improved liquid absorption. Interfering fibers were trimmed with scissors. The residual medium between the tweezer tips was removed by gently inserting a filter paper edge between them. All blotting steps were carried out adjacent to a humidifier to prevent over-drying. Grids were plunged into liquid ethane and stored in liquid nitrogen until imaging. These adjustments reproducibly increased the fraction of grids with suitable ice thickness for tomography (see the histogram in Figure S5). In the absence of the additional front-side blotting step, over 50% of the prepared grids exhibited excessively thick ice, rendering them unsuitable for any type of cryo-EM data acquisition.

Tilt series were collected on a 300 kV Titan Krios G4 (Thermo Fisher Scientific), equipped with a cold field emission gun, a Falcon 4i detector, and a Selectris energy filter set to a 20-eV slit. Data acquisition was conducted in electron-counting mode using tomo5 software (Thermo Fisher Scientific), with the detector gain reference acquired prior to data collection. The tilt range was −60° to +60° in 2° increments. Each tilt image was acquired with a dose of ∼1-1.5 e^-^/Å^2^. Tilt series were recorded at 26,000x magnification (4.64 Å/pixel with a defocus of −10 μm. Tomogram reconstruction was performed using the Etomo graphical interface within the IMOD software package^62^. Tilt series were initially aligned using fiducial-based tracking, followed by fine alignment correction using linear models to minimize residual misalignment. The aligned stacks were reconstructed into tomograms using the Simultaneous Iterative Reconstruction Technique (SIRT) algorithm with 6-8 iterations, depending on sample quality and thickness. Final volumes were binned two times as needed for visualization.

## Competing interests

The authors declare no relevant financial or non-financial interests.

## Author contributions

I.R. and B.Z. designed the research; I.R., J.L., and G.O. performed the experiments, analyzed data, and wrote the manuscript; B.Z. edited the manuscript; T.N. contributed unpublished reagents and analytic tools, and reviewed the manuscript; B.Z. acquired funding. All authors read and approved the final manuscript.

## Acknowledgments

This work was supported by grants from the Swiss National Science Foundation (31003A_179520, 32NE30_185536, CRSII-222809 to BZ). We thank the Dubochet Center for Imaging Bern (DCI-Bern) and the Microscopy Imaging Center (MIC) of the University of Bern for their technical support and access to advanced imaging facilities. We are grateful to Marek Kaminek and David Kalbermatter for their valuable assistance with electron microscopy, and to Beat Haenni and Desirée Dürr for their expert preparation of conventional EM samples. We also thank Diego Haene for his helpful contributions to the experimental work.

## Notes

### Competing Interest Statement

The authors have declared no competing interest.

