## Supplemental table, Supplemental figures for "Advancing Human iPSC-Derived Motor Neuron Models Using Glutamatergic Modulators for Synaptic Function Studies"

### Supplementary Materials

#### *Differentiation conditions*

*Table S1:* ***Overview of experimental timeline and sample collection for functional and structural analyses.*** *This schematic summarizes the differentiation timeline of iPSC-derived motor neurons (iMNs) and indicates the time points at which specific supplements were added. Differentiation days are grouped into defined intervals (DPD0, DPD2–7, DPD7–9, etc.) to highlight the temporal distribution of experiments. iPSCs (yellow) and pre-differentiation stages, EB differentiation stages (orange) and iMN differentiation (pink) reflect progressive neuronal maturation.*

| **Cell stage** | **iPSC** | **EB** | | | | | **iMN** | |
|---|---|---|---|---|---|---|---|---|
| Supplement | DPD0 | DPD0 | DPD 2-7 | D7-9 | D9-14 | D14 | DPD1-14 | DPD14-28 |
| Y-27621 | x | x | | | | | | |
| b-FGF | | x | | | | | | |
| N2 | | x | x | x | x | x | x | x |
| B27 | | x | x | x | x | x | x | x |
| SB431542 | | x | x | | | | | |
| CHIR99021 | | x | x | | | | | |
| LDN193189 | | x | x | | | | | |
| AA | | x | x | x | x | x | x | x |
| RA | | | x | x | x | x | x | x |
| SAG | | | x | x | x | x | | |
| DAPT | | | | | x | x | | |
| BDNF | | | | x | x | x | x | x |
| GDNF | | | | | | x | x | x |
| CNTF | | | | | | x | x | x |
| IGF | | | | | | | x | x |
| CXC156 | | | | | | | | x |
| DPBBD | | | | | | | | x |

*Table S2:* ***Antibodies and reagents used for immunofluorescence, Western blotting and live-cell staining.*** *This table summarizes all primary and secondary antibodies, conjugated antibodies, dyes, and quenchers utilized in the study. For each antibody, the descriptor, manufacturer, product number, host species, target reactivity (species cross-reactivity), and working dilutions for immunofluorescence (IF) and Western blotting (WB) are listed.*

| **Type** | **Descriptor** | **Manufacturer** | **Product number** | **Host** | **Reactivity** | **Dilutions IF/WB** |
|---|---|---|---|---|---|---|
| **Primary Antibodies** | Anti-Synaptophysin | abcam | ab32127 | Rabbit monoclonal | Human, Mouse, Rat | 1:1000/ 1:5000 |
| | Anti-Synapsin | Sigma Aldrich | S193 | Rabbit polyclonal | Human, Mouse, Rat | 1:1000/ 1:5000 |
| | Anti-Rab3a | Proteintech | 16865-1-AP | Rabbit polyclonal | Human, Mouse, Rat | 1:1000/ 1:5000 |
| | Anti-SV2 | abcam | ab77177 | Goat polyclonal | Human, Rat | 1:500/ 1:2000 |
| | Anti-vGlut | Synaptic Systems | 135511 | Mouse monoclonal | Rat, Mouse | 1:1000/ 1:5000 |
| | Anti-ChAT | Invitrogen | MA5-31383 | Mouse monoclonal | Human, Mouse, Rat | 1:1000/ 1:500 |
| | Anti-vAChT | Invitrogen | MA-27662 | Mouse monoclonal | Human, Mouse, Rat | 1:100/ 1:1000 |
| | Anti-beta-III-Tubulin | Novus Biological | NB100-1612 | Chicken polyclonal | Human, Rat | 1:1000/ 1:3000 |
| | Anti-PSD-95 | Synaptic Systems | 124014 | Gunea pig polyclonal | Rat, Mouse | 1: 500/ 1:500 |
| | Anti-PSD-95 | abcam | ab18258 | Rabbit polyclonal | Human, Mouse, Rat | 1:1000/ 1:1000 |
| | Anti-GABAAa1 | Sigma Aldrich | ZRB1626 | Rabbit monoclonal | Human, Mouse, Rat | 1:100/1:1000 |
| | Anti- GluR2 | Sigma Aldrich | ZRB1008 | Rabbit monoclonal | Human, Mouse, Rat | 1:100/1:1000 |
| | Anti-NMDAR | Sigma Aldrich | SAB4501301 | Rabbit polyclonal | Human, Mouse, Rat | 1:1000/ 1:1000 |
| | Anti-GlyR | Synaptic Systems | 146008 | Rabbit monoclonal | Human, Mouse, Rat | 1:500/ 1:1000 |
| | Anti-HB9 | Invitrogen | PA5-23407 | Rabbit polyclonal | Human, Mouse | 1:100/ 1:100 |
| | Anti-islet1 | R&D Systems | AF1837 | Goat polyclonal | Human | 1:100/ 1:100 |
| **Secondary Antibodies** | Anti-Rabbit Alexa-488 | Invitrogen | A11008 | Goat polyclonal | Rabbit | 1:500 |
| | Anti-Rabbit Alexa-568 | Invitrogen | A11011 | Goat polyclonal | Rabbit | 1:500 |
| | Anti-Mouse Alexa-488 | Invitrogen | A11001 | Goat polyclonal | Mouse | 1:500 |
| | Anti-Mouse Alexa-568 | Invitrogen | A11004 | Goat polyclonal | Mouse | 1:500 |
| | Anti-Chicken Alexa-647 | Jackson ImmunoResearch | A16054 | Donkey polyclonal | Chicken | 1:500 |
| | Anti-Guniea pig Alexa-488 | Invitrogen | A11073 | Goat polyclonal | Guniea pig | 1.500 |
| | Anti-Goat Alexa-488 | Invitrogen | A11055 | Donkey polyclonal | Goat | 1:500 |
| | Anti-Goat Alexa-647 | Invitrogen | A21447 | Donkey polyclonal | Goat | 1:500 |
| | Anti-Rabbit HRP | Invitrogen | 31466 | Goat polyclonal | Rabbit | 1:10'000 |
| | Anti-Mouse HRP | abcam | ab6728 | Rabbit polyclonal | Mouse | 1:10'000 |
| | Anti-Chicken HRP | Thermo Fisher Scientific | A16054 | Goat polyclonal | Chicken | 1:10'000 |
| | Anti-Goat HRP | Thermo Fisher Scientific | A15999 | Donkey polyclonal | Goat | 1:10'000 |
| | Anti-Guniea pig HRP | Thermo Fisher Scientific | A18769 | Goat polyclonal | Guniea pig | 1:10'000 |
| | beta III Tubulin HRP | Invitrogen | MA5-16308-HRP | Mouse monoclonal | Human, Mouse etc. | 1:10'000 |
| | GADPH HRP | Invitrogen | MA5-15738-HRP | Mouse monoclonal | Human, Mouse etc. | 1:10'000 |
| **Conjugated Antibodies** | Anti-GluR1 Alexa-647 | Santa Cruz Biotechnology | sc-55509 | Mouse monoclonal | Human, Mouse, Rat | 1:500 |
| | Anti-GluR2 Alexa-568 | Santa Cruz Biotechnology | sc-517265 | Mouse monoclonal | Human, Mouse, Rat | 1:500 |
| **Dyes** | DAPI | Sigma-Aldrich | D9542 | N/A | N/A | 1ug/ml |
| | AM4-64 | Biotium | 70025 | N/A | N/A | 10uM |
| **Quencher** | SCAS | Biotium | 70037 | N/A | N/A | 0.5mM |

*Table S3:* ***Comparison of EM grid types tested for cryo-ET sample preparation of cultured neurons.*** *Cryo-electron tomography (cryo-ET) experiments were conducted on various gold EM grids using standard cell culture methods with minor modifications. The table summarizes each grid type, mesh size, and carbon film configuration, highlighting observed challenges during culture and imaging, adaptations applied to improve performance, and an overall assessment of suitability.*

| Grid type | Mesh No. | Carbon film | Challenges | Adaptation/note | Suitability |
|---|---|---|---|---|---|
| **Quantifoil R2/1, R2/2** | 200 / 300 mesh | Holey carbon | Cells not easily visible; neurites span holes | Suitable for visualization of short neurite | Moderate |
| **Quantifoil S7/2** | 200 mesh | Ultrathin continuous carbon (2–3 nm) | Fragile; unstable for long-term culture | Promotes synapse formation on ultrathin carbon | High |
| **Lacey carbon film** | 200 / 300 mesh | With and without continuous carbon | Cell adhesion and ice thickness issues | Laminin coating improved adhesion | Moderate (most used) |
| **Hexagonal pattern** | Various | With and without continuous carbon | Traps excess media; blotting inefficiency | No viable adaptation found | Unsuitable |

#### *Immunofluorescence of synaptic protein colocalization*

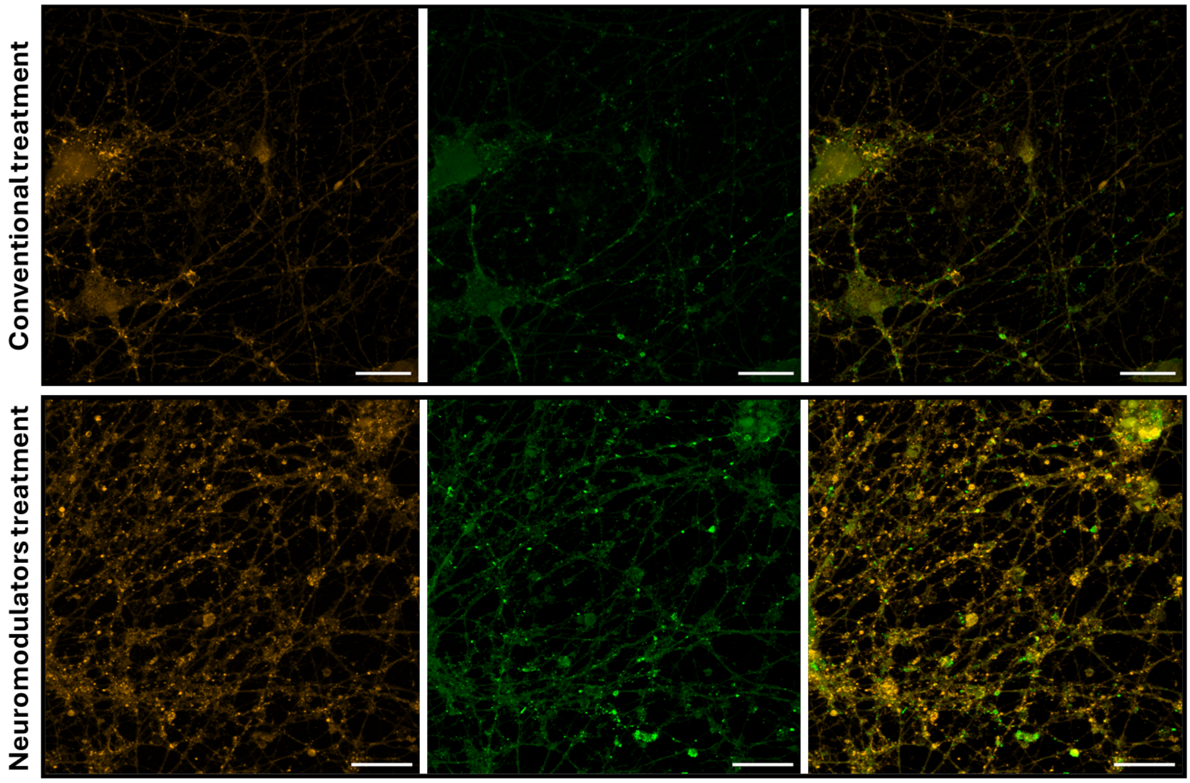


*Figure S1:* ***Confocal analysis of synaptic marker colocalization in iMNs at DPD28 under control and neuromodulator-treated conditions. Top panel (Control):*** *Representative confocal images of untreated iMNs stained for Synaptophysin (SYP, orange), PSD-95 (green), and merged channels. SYP marks presynaptic terminals, while PSD-95 indicates postsynaptic densities. Colocalized puncta along MAP2-positive neurites (not shown here) represent putative excitatory synapses.* ***Bottom panel:*** *iMNs treated with the neuromodulators CX516 and CDPPB exhibit a marked increase in both SYP- and PSD-95-positive puncta, with more extensive colocalization between pre- and postsynaptic markers. This suggests enhanced synaptogenesis following treatment. Scale bars: 20 µm.*

#### *Whole-cell patch clamp electrophysiology analysis*

*Table S4.* ***Quantification of evoked action potential (AP) firing patterns in iMNs.*** Evoked firing properties of induced motor neurons (iMNs) were assessed from whole-cell current-clamp recordings performed at two differentiation stages (DPD21 and DPD28). For each cell, twenty depolarizing current steps (-60 pA to +320 pA, 20 pA increments) were applied. The number of single (sAP) or repetitive (rAP) action potentials was classified during the stimulation phase (SP) and during the subsequent resting phase (RP). Each current injection was considered one independent step.
For each condition (Norm, N; Modulated, M), total event counts were obtained across all recorded cells, separately for rAP SP, sAP SP, rAP RP, and sAP RP categories. The number of recordings (n) and corresponding total steps (exposure = n × 20) were as follows: DPD21 N: 4 coverslips (80 steps), DPD21 M: 4 coverslips (80 steps); DPD28 N: 3 coverslips (60 steps), DPD28 M: 3 coverslips (60 steps). For each category and developmental stage, the relative incidence of events in M compared with N was analyzed using a Poisson-rate model with the exposure term set to the number of steps. To account for multiple testing within each developmental stage (four comparisons per stage: rSP, sSP, rRP, sRP), p-values were adjusted using the Benjamini-Hochberg false discovery rate (FDR) method. Adjusted q-values (qFDR) are reported alongside raw p-values.

***A) DPD21*** *(N: 4 coverslips, 80 steps / M: 4 coverslips, 80 steps)*

| Category | $y_n$ (N) | $y_m$ (M) | RR (M/N) | 95% CI (lower-upper) | Raw p | q-value (FDR) |
|---|---|---|---|---|---|---|
| rAP SP | 24 | 13 | 0.542 | 0.272 - 1.079 | 0.080 | 0.320 |
| sAP SP | 145 | 221 | 1.524 | 1.238 - 1.876 | <0.001 | <0.001 |
| rAP RP | 11 | 12 | 1.091 | 0.491 - 2.425 | 0.830 | 0.996 |
| sAP RP | 23 | 23 | 1.000 | 0.566 - 1.766 | 1.000 | 1.000 |

***B) DPD28*** *(N: 3 coverslips, 60 steps / M: 3 coverslips, 60 steps)*

| Category | $y_n$ (N) | $y_m$ (M) | RR (M/N) | 95% CI (lower-upper) | Raw p | q-value (FDR) |
|---|---|---|---|---|---|---|
| rAP SP | 4 | 14 | 3.500 | 1.157 - 10.587 | 0.029 | 0.116 |
| sAP SP | 128 | 103 | 0.805 | 0.622 - 1.041 | 0.099 | 0.198 |
| rAP RP | 6 | 11 | 1.833 | 0.690 - 4.873 | 0.221 | 0.295 |
| sAP RP | 18 | 10 | 0.556 | 0.258 - 1.195 | 0.134 | 0.268 |

*Table S5.* ***Comparison of spontaneous excitatory postsynaptic currents (sEPSCs) in iMNs under conventional (N) and neuromodulator-enhanced (M) conditions.*** *Paired whole-cell voltage-clamp recordings were analyzed between sister cultures recorded on the same experimental dates under conventional (N) and neuromodulator-treated (M) conditions. Mean values represent per-cell averages. The incidence of spontaneous activity was modeled using a binomial generalized linear model (GLM) with a logit link to estimate the probability of observing at least one sEPSC per recording run. Condition effects were expressed as odds ratios (OR) with 95% confidence intervals (CI). Statistical significance was assessed using two-sided Wald tests, and p-values were adjusted for multiple comparisons using the Benjamini–Hochberg false discovery rate (FDR) method (q-values). Event frequency was modeled using a Poisson GLM with log link and offset for total runs, with condition effects expressed as rate ratios (RR) with 95% CI. The number of recordings (n) and corresponding total runs were as follows: DPD21 N: 23 cells (307 runs), DPD21 M: 29 cells (431 runs); DPD28 N: 10 cells (142 runs), DPD28 M: 15 cells (174 runs).*

***A) DPD21***

| Metric | Ratio (M/N) | 95% CI (lower-upper) | Raw p | q (FDR) |
|---|---|---|---|---|
| Incidence | OR = 1.31 | 0.56 - 3.03 | 0.533 | 0.533 |
| Frequency | RR = 2.30 | 1.28 - 4.14 | 0.005 | 0.007 |

***B) DPD28***

| Metric | Ratio (M/N) | 95% CI (lower-upper) | Raw p | q (FDR) |
|---|---|---|---|---|
| Incidence | OR = 5.48 | 2.77 - 10.83 | <0.001 | <0.001 |
| Frequency | RR = 2.51 | 1.72 - 3.65 | <0.001 | <0.001 |

#### Live staining analysis using AM4-64

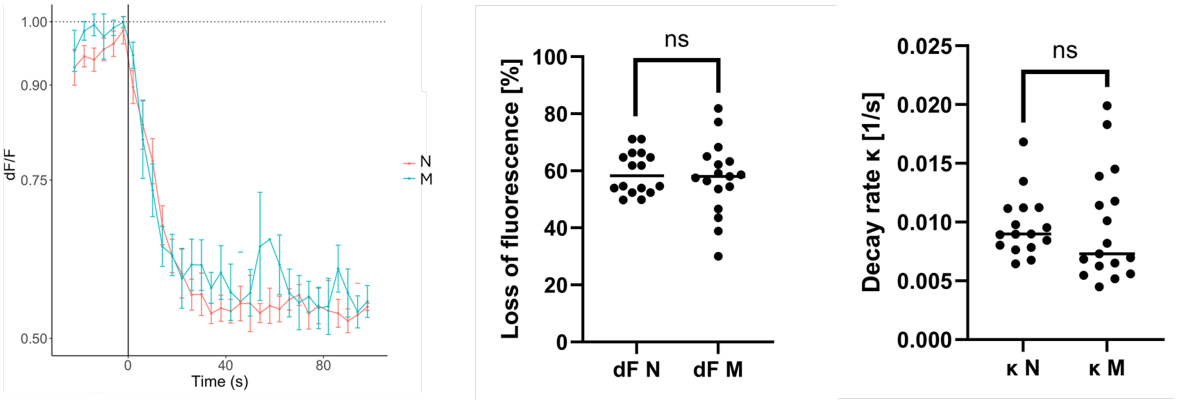


*Figure S2:* ***AM4-64 fluorescence destaining kinetics in human iMNs with or without synapse-modulating treatment. Left panel:*** *Representative fluorescence decay curves (dF/F) of AM4-64-labeled synaptic puncta following high-potassium stimulation (time = 0 s) in iMNs cultured without (orange, N) or with (light blue, M) the neuromodulators CX516 and CDPPB. Both conditions show a rapid drop in fluorescence intensity after stimulation, reflecting SV exocytosis, followed by a gradual plateau phase. Despite a previously observed increase in the density of active puncta in modulator-treated cultures, the overall kinetics of fluorescence decay remain comparable between groups. Error bars represent 95% CI.* ***Middle panel:*** *Quantification of relative loss of fluorescence (%) per punctum, representing the relative fraction of SVs that underwent exocytosis during stimulation. No significant difference was observed between untreated and treated neurons (p-value = 0.58).* ***Right panel:*** *Decay rate constant (κ, in $s^{-1}$) derived from exponential fitting of the fluorescence traces. This parameter reflects the rate of vesicle release (higher κ = faster turnover). No significant differences were detected between conditions (p-value = 0.92; ns = not significant).*

#### Conventional EM of ultrastructures in iMNs

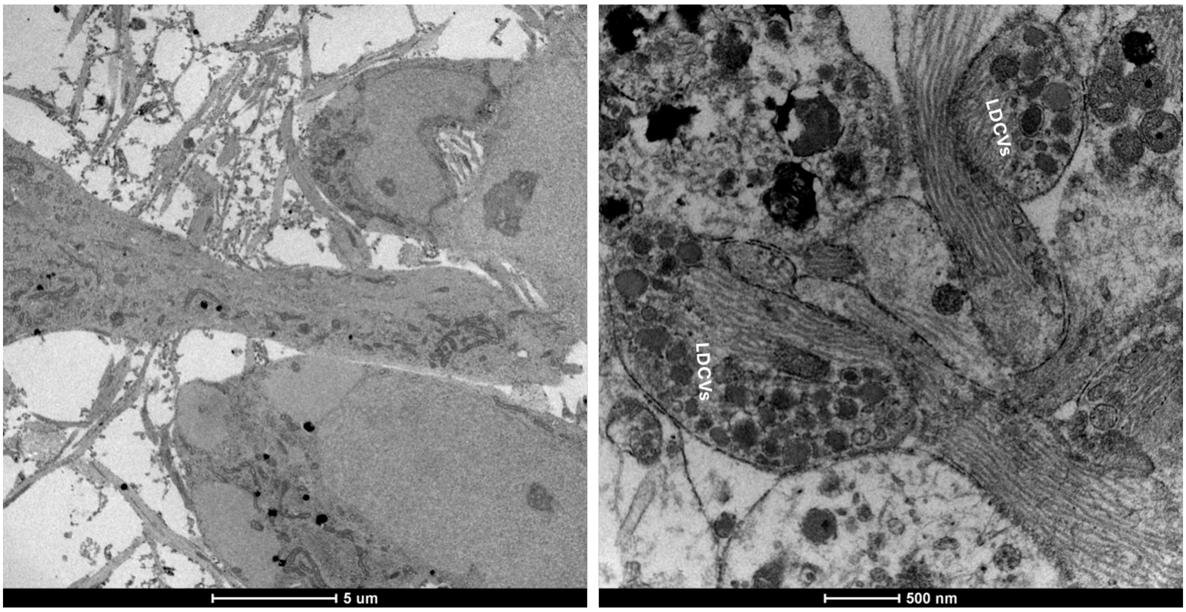


*Figure S3:* ***Electron micrographs of selected iMN characteristics. Left micrograph:*** *An overview of healthy neurons at DPD28 with presence of neurites surrounding the somata.* ***Right micrograph:*** *Large dense core vesicles (LDCVs) (50-100 nm diameter) localized in varicosities.*

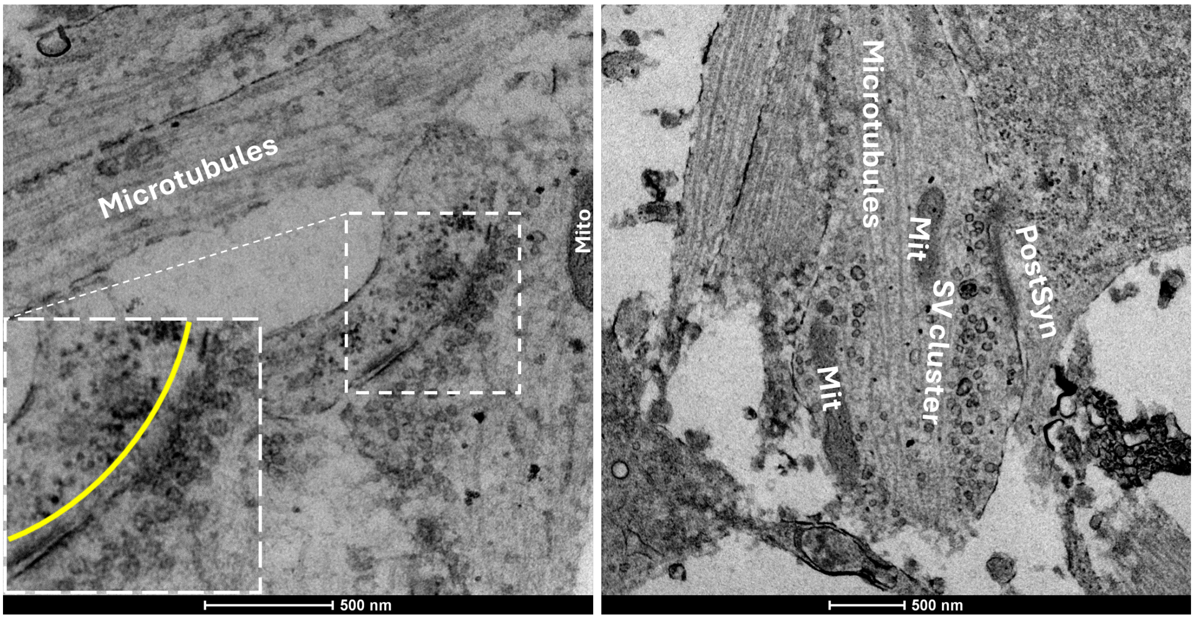


*Figure S4:* ***Ultrastructural features of synaptic terminals in iMNs following neuromodulator treatment, revealed by conventional transmission EM. Left panel:*** *Representative EM micrograph showing a large presynaptic bouton with a densely packed SV cluster adjacent to the presynaptic membrane, indicative of a prominent active zone. Numerous vesicles appear docked, suggesting an expanded readily releasable pool consistent with enhanced synaptic activity following treatment with CX516 and CDPPB. Inset shows a magnified view of the active zone region. The yellow curved line marks the postsynaptic zone, which may represent one large or two closely apposed postsynaptic compartments.* ***Right panel:*** *EM image of an* en passant *synapse highlighting key ultrastructural components of a mature neuronal terminal. Microtubules are aligned along the axon shaft, while mitochondria (Mit) are positioned near the vesicle cluster. The presynaptic terminal contains a tightly organized SV cluster, aligned with the postsynaptic membrane (PostSyn), demarcating a clearly identifiable synaptic cleft.*

#### *Cryo-EM/ET sample preparation; plunge freezing the grids*

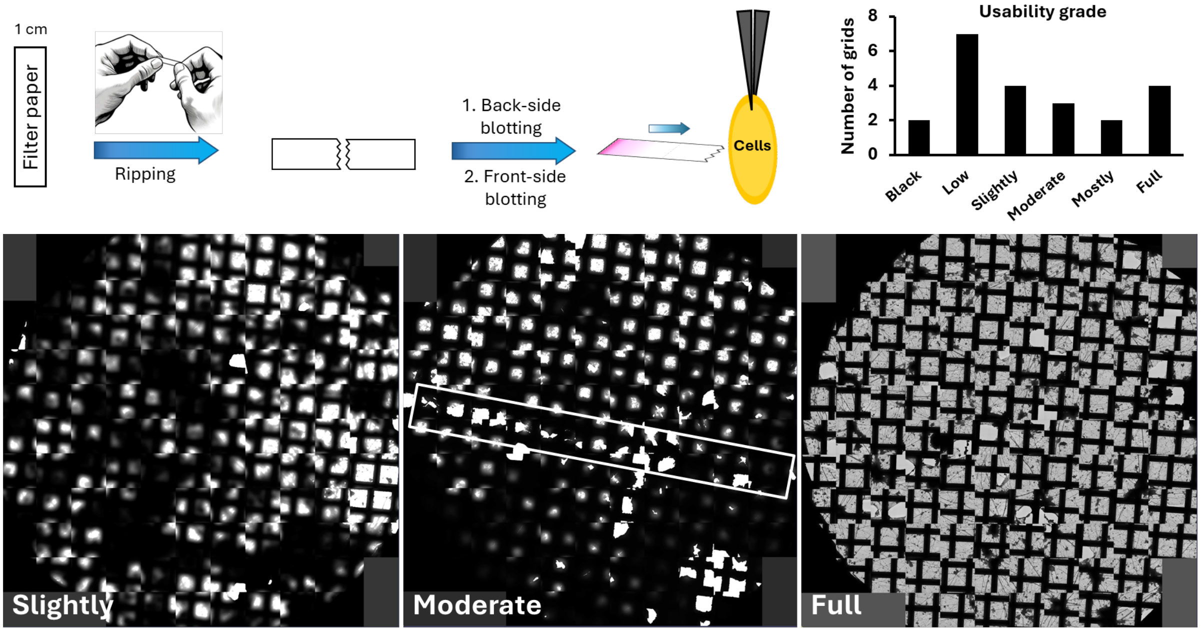


*Figure S5:* ***Schematic of the front-blotting procedure using a torn filter paper edge and assessment of cryo-EM grid usability following front-side blotting optimization.*** *To reduce ice thickness and improve grid quality for cryo-electron tomography, an additional manual front-side blotting step was introduced: 1) A standard filter paper is torn by hand to create a jagged edge with irregular protrusions that enhances the capability of liquid absorbance. 2) This torn edge is used to gently blot the front side of the EM grid near the area containing the cell monolayer. 3) Blotting is performed perpendicular to the grid surface and closer to the tweezers, targeting the periphery without disturbing the central cell*

area. The histogram on the right shows the evaluation of 22 cryo-EM grids prepared using the modified front-side blotting approach with torn filter paper edges. Each grid was assessed for vitrification of quality and suitability for cryo-ET and scored based on the proportion of usable area suitable for data acquisition. The histogram displays the distribution of usability grades across all tested grids. Notably, most of the grids were usable for image acquisition. This demonstrates the effectiveness of the front-side blotting method in consistently producing grids with proper ice thickness. ***Bottom panel*** *shows representative examples of intuitive usability grades assigned to grids based on stitched low-magnification overviews. The white rectangle outlines a potential trace of the blotting edge, supporting the directional effect of perpendicular front-side blotting.*

#### *Cryo-ET of ultrastructures in iMNs*

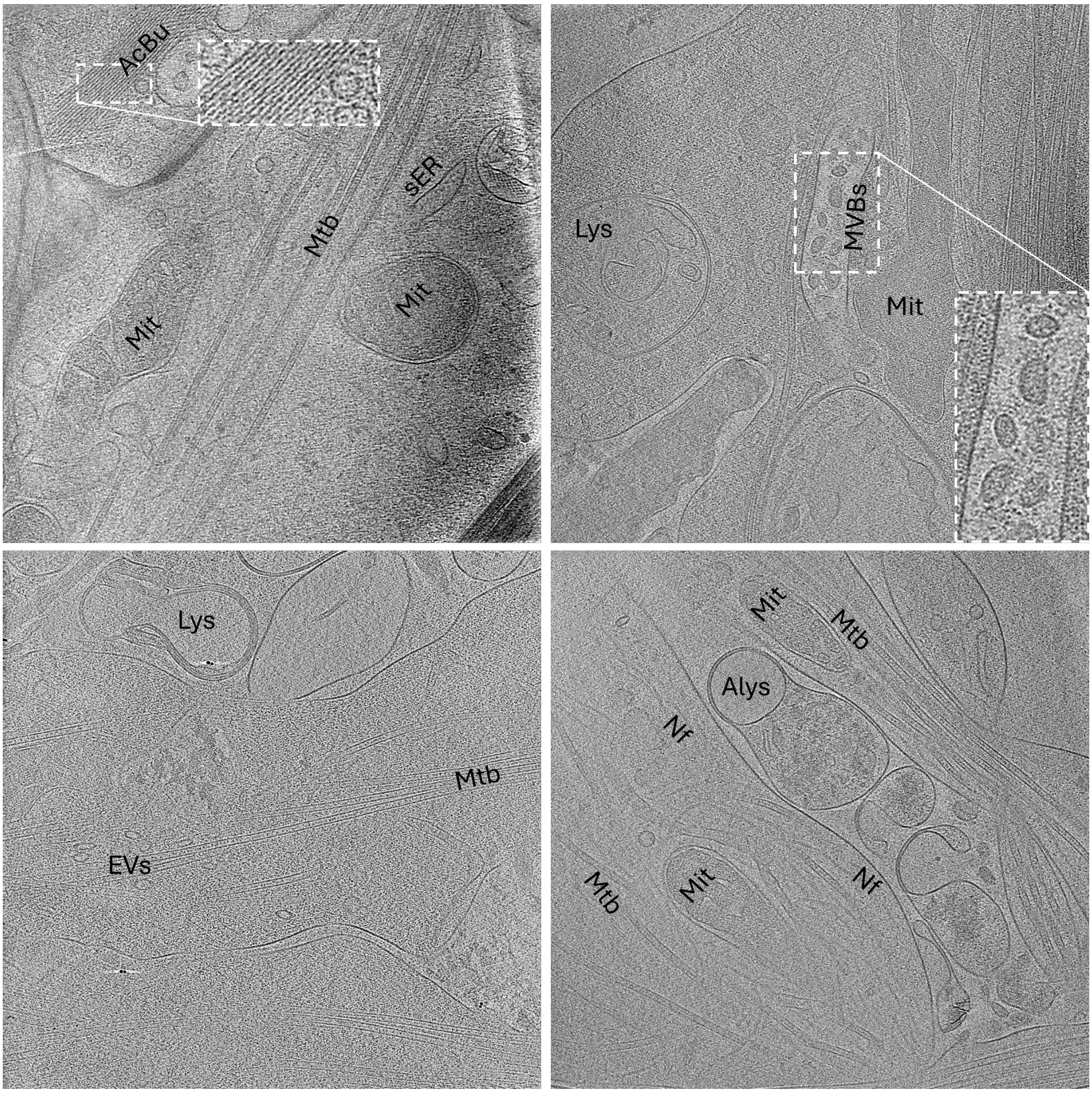


*Figure S6:* ***Cryo-ET reveals the ultrastructural organization of cytoskeletal and organelle networks in neuronal extensions.*** *Representative tomographic slices from axonal and dendritic regions of iMNs show a well-organized cytoskeletal architecture. Parallel bundles of microtubules (Mtb) serve as structural tracks for intracellular transport, interspersed with elliptical vesicles (EVs) likely in transit. Neurofilaments (Nf) and actin filaments (AcBu) provide additional mechanical support and remodeling flexibility. Smooth Endoplasmic reticulum (sER) tubules are distributed throughout the cytoplasm and neurites, often near mitochondria (Mit) with clearly defined cristae. Multivesicular bodies (MVBs) and various stages of lysosomal compartments, including lysosomes (Lys) and autolysosomes (Alys), reflect active degradation and membrane trafficking. The presence of extracellular Lys and Alys may result from either plasma membrane fusion events or cellular lysis. This integrated framework highlights the complexity of intracellular transport, organelle distribution, and membrane dynamics in neuronal processes.*

#### Synaptic activity-dependent endocytosis revealed by ferritin uptake in iMNs

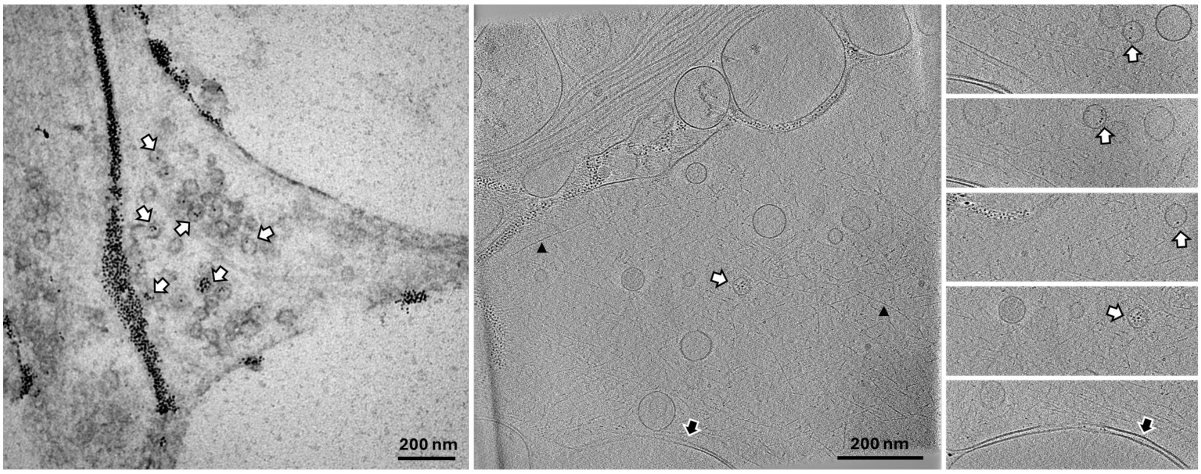


*Figure S7:* ***Visualization of synaptic activity-dependent endocytosis in iMNs using cationized ferritin uptake.*** *Transmission electron micrograph showing a neurite of an iMN following exposure to cationized ferritin and high-potassium stimulation. Electron-dense ferritin particles are visible within small vesicles and endosome-like compartments (50–200 nm in diameter), clustered near the plasma membrane. This labeling indicates successful internalization via activity-dependent endocytosis. The dense accumulation of labeled vesicle-like structures reflects active synaptic vesicle recycling.* ***Left panel*** *shows conventional EM;* ***middle panel*** *shows cryo-ET tomographic slice with the montages from different planes of the tomogram on the* ***right panel****. White filled arrows show the recycled vesicles containing ferritin particles, black filled arrow shows the possible synaptic connection with a gap of 20 nm, and black arrow heads show the actin filaments.*

#### Dose-dependent neurotoxicity of glutamatergic modulators in iMNs

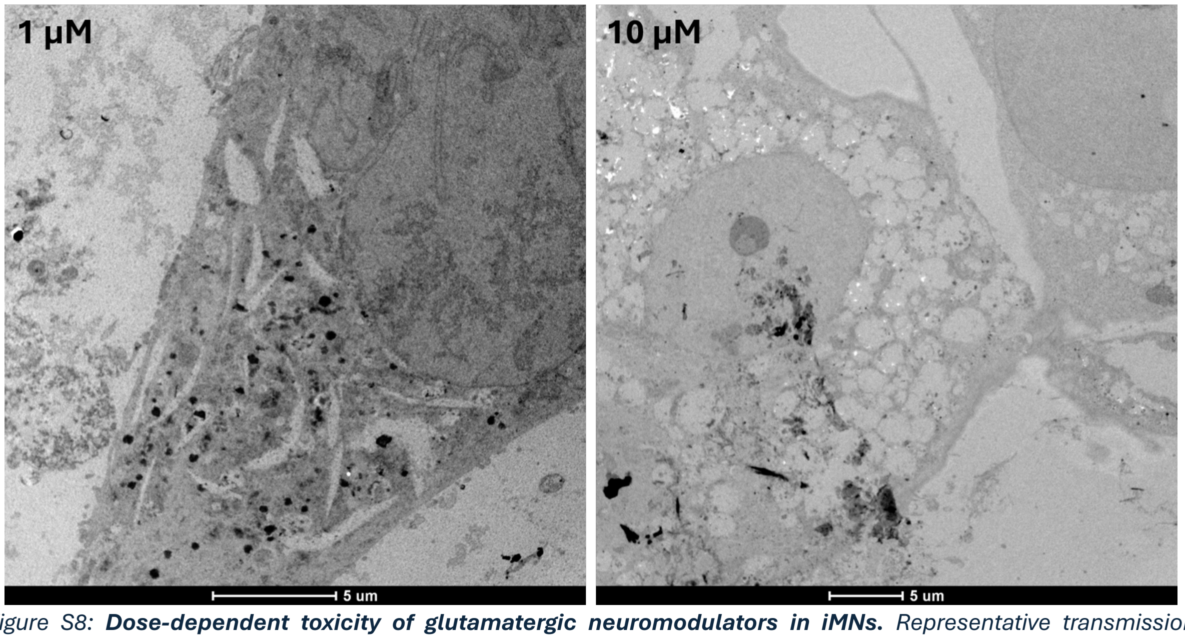


*Figure S8:* ***Dose-dependent toxicity of glutamatergic neuromodulators in iMNs.*** *Representative transmission electron micrographs of iMNs treated with CX516 and CDPPB at 1 µM (left) and 10 µM (right), one week after addition at DPD14. While neurons exposed to 1 µM show preserved subcellular architecture with moderate vacuolization, cells treated with 10 µM exhibit widespread cytoplasmic disintegration, vacuole accumulation, and organelle degradation, indicative of pronounced neurotoxicity. These findings informed us of the choice of 100 nM as the optimal working concentration in all functional experiments.*

#### *Live calcium imaging of iMNs under baseline and depolarizing conditions*

*Movie S1a:* ***Spontaneous calcium activity in iMNs.*** *Time-lapse video of iMNs under baseline HEPES ACSF conditions (no external trigger), showing spontaneous Fluo 4 fluorescence fluctuations indicative of intrinsic network activity. Frame interval: 3.22 s (10 fps).*

*Movie S1b:* ***Depolarization-evoked calcium influx in iMNs.*** *Time-lapse video recorded before, during, and after application of high $K^+$ HEPES ACSF (50 mM $K^+$). The video shows a robust, synchronized calcium influx across neuronal somata and processes, followed by gradual recovery during washout. Frame interval: 3.22 s (10 fps).*
